# Stromal DUF760-1 and DUF760-2 proteins are degraded by the chloroplast Clp protease system but have very different half-lives

**DOI:** 10.1101/2024.03.10.584298

**Authors:** Bingjian Yuan, Klaas J. van Wijk

## Abstract

The chloroplast AAA+ chaperone CLPC1 aids to select, unfold and deliver hundreds of proteins to the stromal CLP protease core complex for degradation. Through *in vivo* CLPC1 trapping we previously identified dozens of trapped proteins that are substrate adaptors (*e.g.* CLPS1 and CLPF), other chaperones (CLPC2 and CLPD) or (potential) substrates (*e.g.* RH3) for the CLP chaperone-protease system. Here we show that two of these highly trapped proteins, DUF760-1 and DUF760-2, are substrates for the chloroplast CLP protease system. *In planta* BiFC and yeast-2-hybrid analyses show that both DUF760 proteins can directly interact with the N-domain of CLPC1. Immunoblotting and confocal microscopy analysis demonstrates that both DUF760 proteins are highly enriched in *clpc1-1* and *clpr2-1* loss-of function mutants. *In vivo* cycloheximide chase assays in different genetic backgrounds show that DUF760-1 and 2 are both degraded by the chloroplast CLP protease. The half-life of DUF760-1 is 4-6 hours, whereas DUF760-2 is so unstable that it is very hard to detect unless degradation is inhibited in CLP loss-of-function alleles. Null mutants for DUF760-1 and DUF760-2 show weak but differential pigment phenotypes and differential sensitivity of protein translation inhibitors. The functions of these DUF760 proteins are unknown; the lack of shared mRNA co-expressors and the large difference in half-life and protein abundance, suggest that they play different roles within the chloroplast. Taken together, our results demonstrate that DUF760-1 and 2 are newly discovered substrates of CLP chaperone-protease system.

## INTRODUCTION

Intra-chloroplast proteolysis is a key part of chloroplast proteostasis and plays a crucial role in the development and function of chloroplasts (Gao et al., 2023; Llamas and Pulido, 2022; Majsec et al., 2017; Nishimura et al., 2016). Chloroplasts contain several dozen different proteases organized as either homomeric proteins or organized into heteromeric complexes. These proteases are distributed across different intra-chloroplast compartments, including the thylakoid lumen (*e.g.* CTPA, DEG1,5,8), the thylakoid membrane (*<u>e.g</u>*. FTSH1,2,5,8, EGY1,2, SPPA), plastoglobules (PGM48) the chloroplast stroma (*e.g.* CLP, CGEP, SPP, PREP1,2, OOP, various aminopeptidases) as well as the inner envelope membrane (*e.g.* FTSH7,8,9,11,12 and rhomboids) (Gao *et al*., 2023; Llamas and Pulido, 2022; Majsec *et al*., 2017; Nishimura *et al*., 2016). Intra-chloroplast proteolysis is complemented by chlorophagy and other extra-chloroplast proteolytic pathways in which chloroplast proteins are degraded in the cytosol or vacuole (Fu et al., 2022; Izumi and Nakamura, 2018; Woodson, 2022).

Within the intra-chloroplast proteolysis network, the soluble stromal CLP chaperone-protease system plays a key role in the degradation likely of hundreds of proteins (Bouchnak and van Wijk, 2021; Nishimura and van Wijk, 2015; Rodriguez-Concepcion et al., 2019). The chloroplast CLP protease consists of a tetradecameric CLPPR protease core complex formed by two heptameric rings (the P-ring with CLPP3-6 and the R-ring with CLPP1 and CLPR2-4) and the CLPT1 and CLPT2 proteins (Bouchnak and van Wijk, 2021). Three AAA+ chaperones (CLPC1, CLPC2, and CLPD) form hexamers that recognize substrates through their N-domains, and these hexamers unfold and deliver substrates into the CLPPRT core complex driven by ATP hydrolysis. The adapters (CLPS1 and CLPF) function in substrate recognition and delivery to CLPC (Bouchnak and van Wijk, 2021).

Several studies have shown that chloroplast CLPC1 and the CLP protease are required for the degradation of the thylakoid P1B-type ATPase PAA2/HMA8 (Tapken et al., 2015), deoxyxylulose 5-phoshate (DXS) (Pulido et al., 2016), phytoene synthase (PSY) (Welsch et al., 2018), genomes uncoupled 1 (GUN1) (Wu et al., 2018), and glutamyl-tRNA reductase (GluTR) (Apitz et al., 2016; Nishimura et al., 2015; Richter et al., 2019). CLPS1 affinity studies identified a dozen chloroplast stromal targets for this N-recognin, including GluTR and three enzymes within the aromatic amino acid biosynthesis (DAHP, DHS, CS) (Nishimura et al., 2013). Comparative proteomics studies of several CLP mutants identified many additional candidate substrates as well as altered protein accumulation due to indirect effects of reduced CLP capacity such as upregulation of chloroplast chaperones; however, these studies cannot distinguish between direct and indirect effects of reduced CLP capacity (Bouchnak and van Wijk, 2021).. We previously used an *in vivo* trapping approach to identify interactors with CLPC1 in *Arabidopsis thaliana* by expressing a STREPII-tagged copy of CLPC1 mutated in its Walker B domains (CLPC1-TRAP) followed by affinity purification and tandem mass spectrometry (MSMS) (Montandon et al., 2019; Rei Liao et al., 2022). These trapping studies identified 59 highly enriched CLPC1 interacting proteins involved in chloroplast metabolism as well as proteins belonging to families of unknown functions (DUF760, DUF179, DUF3143, UVR-DUF151, HugZ/DUF2470), as well as the UVR domain proteins EXE1 and EXE2 implicated in singlet oxygen damage and retrograde signaling (Dogra et al., 2019; Dogra et al., 2021). Among these trapped proteins were DUF760-1 (AT1G32160) and DUF760-2 (AT1G48450).

In this study, we investigated the relation between DUF760-1,2 and the chloroplast CLP chaperone-protease system and we demonstrated that both proteins are novel substrates of the CLP protease system. DUF760-1,2 can interact with the N-domain of CLPC1. DUF760-1 and DUF760-2 proteins are multiple fold enriched in the *clpc1* and *clpr2-1* mutants and *in vivo* cycloheximide (CHX) chase experiments showed that the CLP protease system is required for DUF760-1,2 degradation. Interestingly, the half-life of DUF760-2 is extremely short (minutes) explaining why DUF760-2 accumulates at very low levels in wild-type plants, but at high levels in the CLPC1-TRAP line or CLP loss-of-functions mutants. This study also demonstrates that our *in vivo* trapping strategies with a mutated ClpC1 chaperone impaired in ATP hydrolysis (Montandon *et al*., 2019; Rei Liao *et al*., 2022; Rei Liao and van Wijk, 2019) is an excellent and effective tool to identify novel CLP substrates including proteins that normally accumulate at very low steady state level (often not detected by MSMS) due to very high turn-over rates.

## RESULTS

### DUF760-1 and DUF760-2 are plastid proteins with different protein abundances and co-expressors, but similar overall mRNA abundance and distribution

DUF760-1 and DUF760-2 were identified by MSMS in our recent *in vivo* CLPC1 trapping study (Rei Liao *et al*., 2022) DUF760-1 was 4-fold enriched in the CLPC1-TRAP-STREPII eluates compared to CLPC1-WT-STREPII eluates (Table 1). DUF760-2 was identified with an extremely high number of matched MSMS spectra (584) in the CLPC1-TRAP eluates and only 16 in the CLPC1-WT-STREPII eluates; this showed that DUF760-2 was 32-fold enriched in the CLPC1-TRAP-STREPII compared to CLPC1-WT-STREPII (Table 1). Interesting, inspection of the Arabidopsis PeptideAtlas (https://peptideatlas.org/builds/arabidopsis), which reports MSMS-based protein identifications based on ∼70 million matched MSMS spectra collected across 115 different studies (van Wijk et al., 2024), shows that DUF760-1 has a much higher abundance across tissues than DUF760-2 (Table 1). Hence the extremely high abundance of DUF760-2 in the CLPC1-TRAP affinity eluates compared to the CLPC1-WT affinity eluates suggests that DUF760-2 is stabilized in the CLPC1-TRAP background and directly or indirectly interacts with the CLPC1-TRAP chaperone.

**Table 1.** DUF760-1,2 abundance in CLPC1 affinity experiments and in PeptideAtlas.

|  | *in vivo* affinity trapping<br>(adjSPC) (a) |  | PEPTIDE ATLAS (b) |
| --- | --- | --- | --- |
|  | CLPC1-WT-<br>STREPII | CLPC1-TRAP-<br>STREPII | # experiments |
| **DUF760-1** | 4 | 19 | 135 |
| **DUF760-2** | 16 | 584 | 19 |

(a) adjusted spectral counts - from Liao et al (2022) JBC
(b) https://peptideatlas.org/builds/arabidopsis

mRNA expression atlas data in ePlant (http://bar.utoronto.ca/) show that DUF760-1 and DUF760-2 have similar tissue and developmental mRNA expression patterns (Supplemental Figure 1), which is surprising given the much higher protein abundance of DUF760-1. Construction of mRNA-based co-expression networks for the CLP genes and trapped proteins showed that these two *DUF760* genes operate in different parts of the network (Rei Liao *et al*., 2022). A direct comparison of the top 50 co-expressors for DUF760-1 and DUF760-2 from the ATTEDII data base (https://atted.jp/) (Obayashi et al., 2022) show that both genes co-express nearly exclusively with genes encoding for chloroplast proteins (co-expression with genes encoding for other chloroplast proteins is typical (Majsec *et al*., 2017)), but that there is hardly any overlap in co-expressors (Supplemental Table 1). DUF760-1 and DUF760-2 both are predicted to be sorted into the chloroplast and the most N-terminal peptides identified by MS collected from the Arabidopsis PeptideAtlas (van Wijk *et al*., 2024) closely agree with the predicted cTP cleavage site (Supplemental Figure 2), providing strong support for intra-chloroplast accumulation.

### Spatiotemporal accumulation of DUF760-1 and DUF760-2 proteins

To further determine the subcellular localization and spatiotemporal abundance of DUF760-1,2, we generated stable transgenic plants expressing genomic DNA of *DUF760-1* or *DUF760-2* fused to a C-terminal YFP tag in wild-type (WT) plants (*g-DUF760-1/WT* and *g-DUF760-2/WT*) (see Supplemental Table 2 for the nomenclature and information about transgenic lines used in this study). Immunoblot analysis of total cellular protein extracts of different parts of these transgenic lines showed that DUF760-1 accumulated in young rosette leaves, stems, flowers and green siliques, but only very little in roots and older rosettes (Figure 1A). Consistent with these immunoblotting results, confocal microscopy analysis of these transgenic lines detected DUF760-1 protein in cotyledons and rosette leaves (Figure 1A), as well as other green tissues such as hypocotyls, siliques and sepals, but was undetected in root (Supplemental Figure 3A). Immunoblot analysis of the same protein extracts showed that DUF760-2 accumulated only at very low levels in young leaves and in roots and was very hard to detect in other plant tissues (Figure 1B). We note that the same protein extracts and commercial anti-GFP serum were used for DUF760-1 and DUF760-2 which allows for direct comparison of these proteins. Consistent with these immunoblotting results, confocal microscopy analysis of these transgenic lines detected DUF760-2 protein at low levels in cotyledons and young rosette leaves (Figure 1B), and generally very low levels in other green tissues. Interestingly and consistently with the immunoblot, some accumulation was observed by confocal microscopy in roots tips (Supplemental Figure 3B).

**Figure 1.**
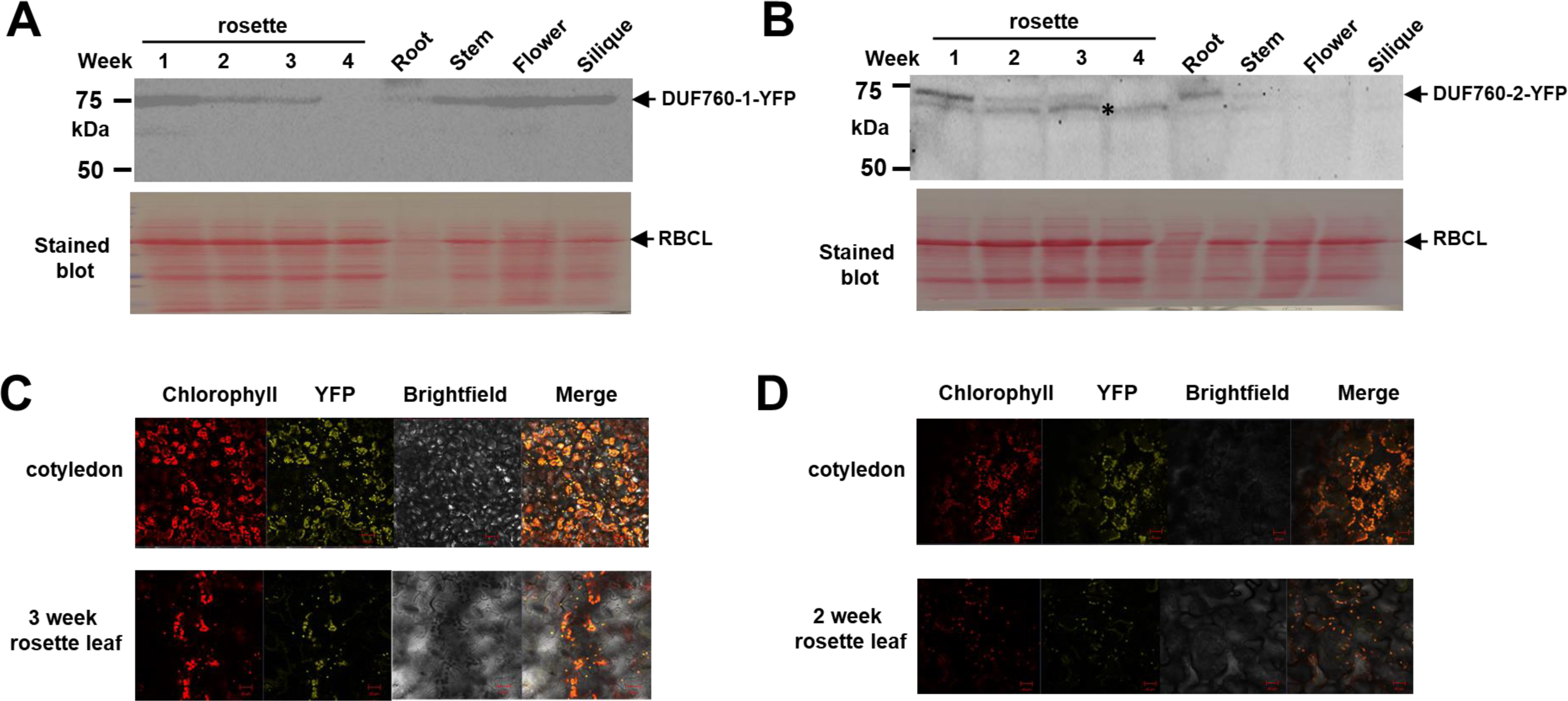
Protein accumulation of DUF760-1 and DUF760-2 in different organs and developmental stages, and subcellular localization in *g-DUF760-1-YFP*/WT and *g-DUF760-2-YFP*/WT. WT, *g-DUF760-1-YFP*/WT and *g-DUF760-2-YFP*/WT plants were grown in controlled growth chambers under constant light of 100 μmol photons.m^-1^.s^-1^ and a relative humidity of 50%–70%. **(A, B)** Accumulation of *Arabidopsis* DUF760-1-YFP **(A)** and DUF760-2-YFP **(B)** in leaves, stems, flowers, siliques, and roots as determined by immunoblotting with α-GFP antibodies. The lower panels show the Ponceau-stained blot prior to incubation with antibodies; the dominant band of RBCL is marked. Asterisk marked an unspecific response to α-GFP. **(C, D)** Confocal microscopy of cotyledon and rosette leaves of *g-DUF760-1-YFP*/WT **(C)** and *g-DUF760-2-YFP*/WT **(D)**. The panels show signals from chlorophyll, YFP, the brightfield image and the merger of the three panels.

### Characterization of *duf760-1,2* null mutants and genetic interactions between DUF760-1,2 and CLPC1

To investigate the biological functions of DUF760-1,2, we searched for publicly available T-DNA lines and obtained two T-DNA insertion lines (*duf760-1-1*, *duf760-1-2*) for *DUF760-1* and one line (*duf760-2-1*) for *DUF760-2* (Figure 2A). RT-PCR analysis showed a complete loss of transcripts of *DUF760-1* and *DUF760-2* in the respective T-DNA lines, indicating that these are all null mutants (Figure 2B). Growth and development of these mutants grown on agar plates (not shown) or soil did not show obvious visible phenotypes compared to WT (Figure 2C). However, chlorophyll and carotenoid accumulation per fresh weight was significantly reduced in *duf760-2-1* but not in *duf760-1-1,2* (Figure 3A, B). Expression of genomic *DUF760-2* in *duf760-2-1* fully complemented this reduced pigment phenotype (Figure 3B).

**Figure 2.**
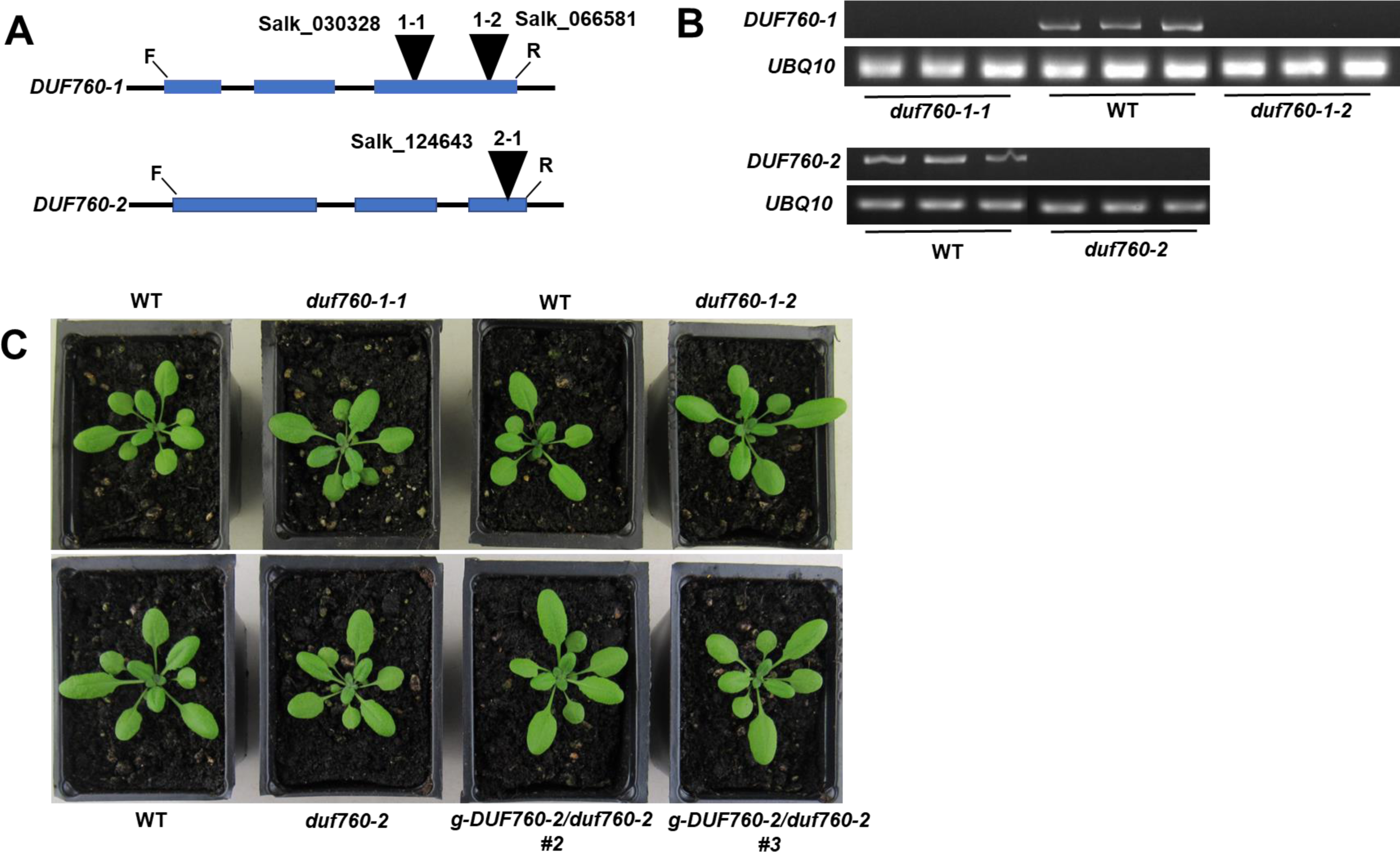
Analysis of null mutants of *DUF760-1* and *2* in *Arabidopsis*. **(A)** Gene model structure and position of T-DNA insertions in the mutants of *DUF760-1* and *DUF760-2* used in this study. Exons (blue boxes), intron (black lines), and T-DNA insertions (triangle) are indicated. **(B)** mRNA levels of *DUF760-1* and *DUF760-2* in the WT, *duf760-1-1,2* and *duf760-2*. Transcripts were amplified by RT-PCR (25 cycles) with gene-specific primers and analyzed on an agarose gel. *UBQ10* mRNA was used as an internal control. **(C)** Phenotypic analysis of the WT, *duf760-1-1,2*, *duf760-2* and two independent *duf760-2* complemented lines. Plants were grown for 20 d on soil under 16/8-h light/dark cycle at ∼100 µmol photons m^−2^ s^−1^. No visible differences were observed.

**Figure 3.**
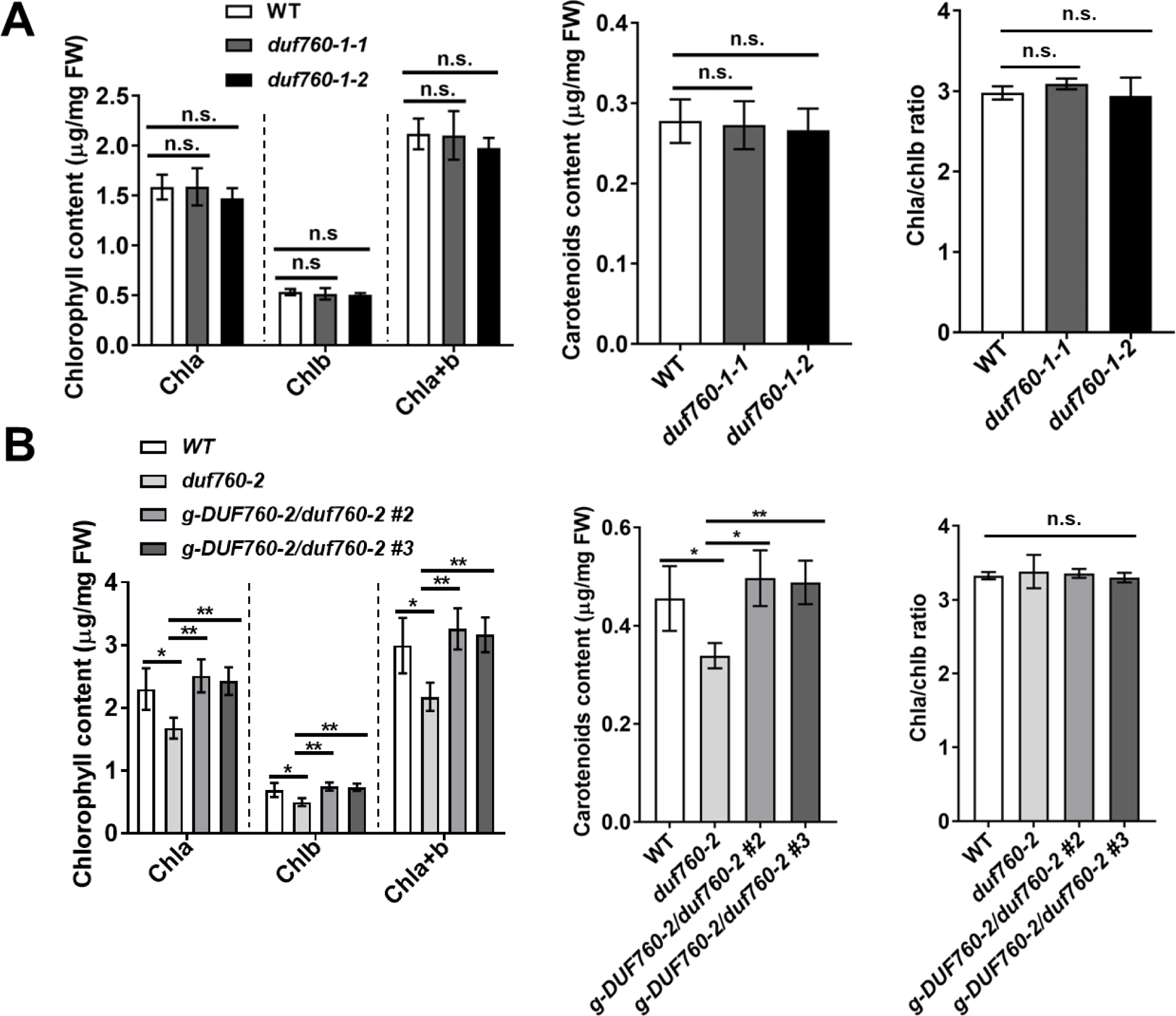
Chlorophyll and carotenoid analysis of *duf760-1-1,2, duf760-2* and *duf760-2* complemented lines at development stage 1.07. **(A, B)** WT and *duf760-1-1,2* **(A)** and WT, *duf760-2* and *duf760-2* complemented lines **(B)** were grown for 20 d on soil under 16/8-h light/dark cycle at ∼100 µmol photons m^−2^ s^−1^. Chlorophyll a+b (Chla+b) and total xanthophyll and other carotenoid (X+C) contents were determined on a fresh weight basis; *n* = 3. SE is indicated. *p ≤ 0.05, **p ≤ 0.01; n.s. - not significant (two-tailed t-test).

The antibiotics CHX and chloramphenicol (CAP) are inhibitors of cytoplasmic and mitochondrial/chloroplast protein synthesis, respectively. Our previous study regarding the adaptor CLPS1 showed that the *clps1* null mutant has an enhanced sensitivity to CAP but not CHX (Nishimura *et al*., 2013). This inspired us to investigate the impact of these translation inhibitors on *duf760-1-1*, *duf760-1-2* and *duf760-2-1*. WT and these three null mutants were grown on MS medium with sucrose supplemented with CAP and CHX for 20 days under short day conditions (Figure 4, Supplemental Figure 4). *duf760-2-1* was significantly more sensitive to both CAP and CHX (p<0.01), and this could be fully complemented by transformation with genomic *DUF760-2* (Figure 4). In contrast, *duf760-1* mutants did not show significant (p<0.01) altered sensitivity to these antibiotics (Supplemental Figure 4). We conclude that the loss of DUF760-2 makes plants more susceptible to proteostasis challenges.

**Figure 4.**
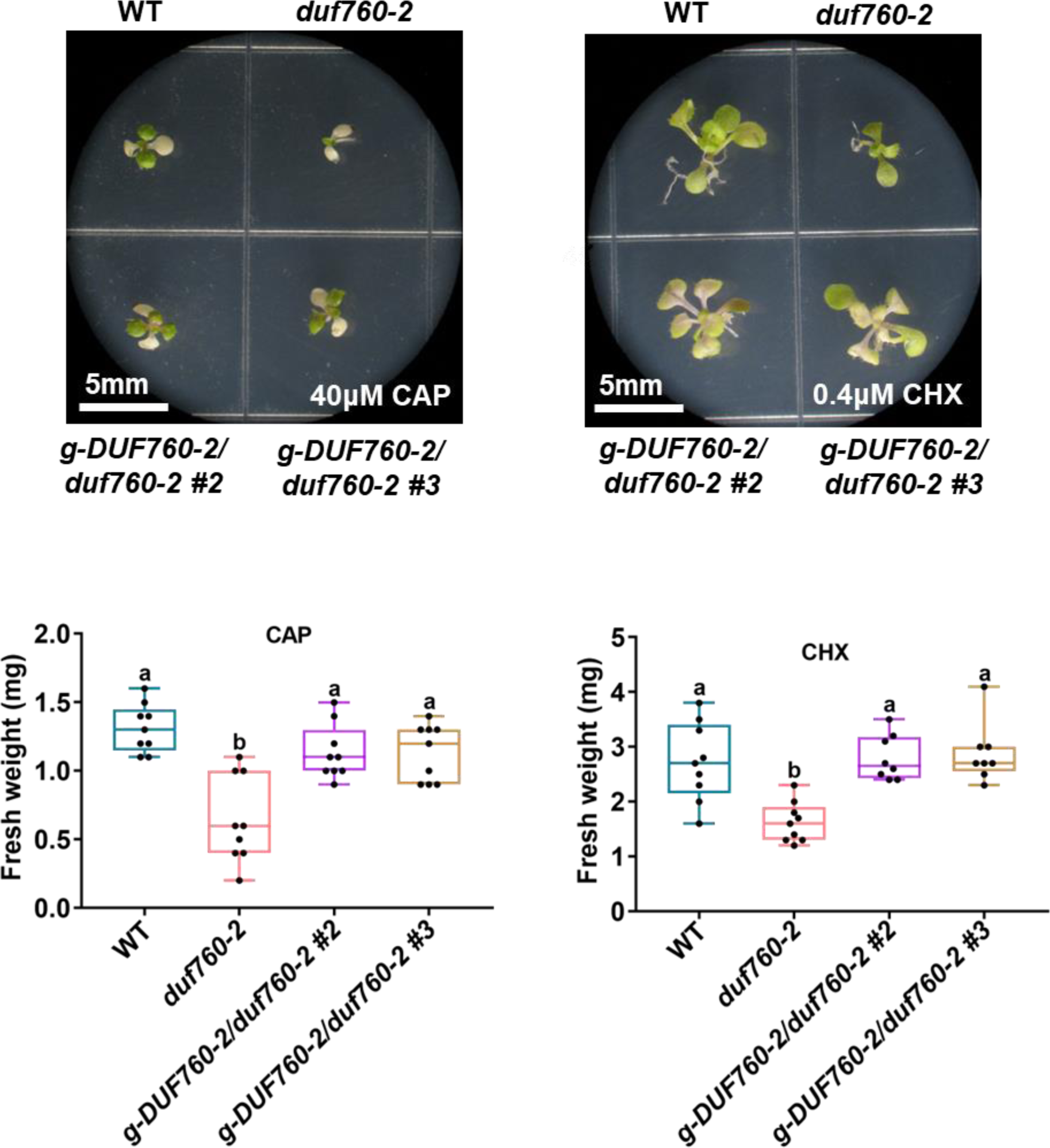
Effects of translation inhibitors on the WT and *duf760-2*. **(A)** Seedling phenotypes of WT, *duf760-2* and complemented *duf760-2* lines after treatment with CAP or CHX. Plants were grown for 20 d under short days on half-strength Murashige and Skoog medium containing 2% sucrose and 40μM CAP or 300 μM CHX. Bars = 5 mm. **(B)** Effect of CAP or CHX treatment on fresh weight of WT and *duf760-2* seedlings. Error bars indicate SD. Lowercase letters indicate significant differences between multiple groups by one-way ANOVA at p<0.01.

We then tested the genetic interactions between *DUF760-1*,*2* and *CLPC1*. Double mutants of *clpc1-1 duf760-1-1* and *clpc1-1 duf760-2* were generated by crossing and screening by genotyping of the resulting F2 and F3 progenies for homozygous double null mutants (Supplemental Figure 5). *clpc1-1 duf760-1-1* and *clpc1-1 duf760-2* showed the same virescent and reduced growth phenotype as *clpc1-1*. We also created the *duf760-1 duf760-2* double mutants by crossing, but we did not observe any genetic interactions (data not shown).

In addition, to further investigate the functions of *DUF760-1* and *2* in *Arabidopsis* chloroplast, we generated the overexpression transgenic lines of *DUF760-1* and *2* by expressing *35S-DUF760-1-GFP* or *35S-DUF760-2-GFP* in WT (*35S-DUF760-1*/WT and *35S-DUF760-2*/WT). Two independent transgenic lines for each gene were confirmed by RT-PCR assays and selected for further study (Supplemental Figure 6A and 6B). Growth and development of homozygous *35S-DUF760-1*/WT and *35S-DUF760-2*/WT when grown on soil were not visibly different from the wild type (Supplemental Figure 6A). In addition, the chlorophyll and carotenoid contents of *35S-DUF760-1*/WT and *35S-DUF760-2*/WT are comparable to those in WT (Supplemental Figure 6C).

### DUF760-1 and DUF760-2 over-accumulate in *clpc1-1* and *clpr2-1*

Multiple studies have shown that the substrates of CLP proteases (*e.g.* PAA2, DXS, GluTR, PSY, GUN1) over-accumulate in various *clp* mutants (Apitz *et al*., 2016; Nishimura *et al*., 2015; Pulido *et al*., 2016; Richter *et al*., 2019; Welsch *et al*., 2018). Thus, we hypothesized that if DUF760-1 and DUF760-2 are targets of the CLP protease system, both DUF proteins are expected to be elevated in *clp* mutants, such as *clpc1-1* and *clpr2-1*. To test this possibility, we introduced *35S-DUF760-1-GFP* or *g-DUF760-2-YFP* in the mutants of *clpc1-1* and *clpr2-1* by crossing; genotypes of F2 progenies were confirmed by genotyping and RT-PCR (Supplemental Table 2; Figures 9 and 6).

The transgenic plants of *35S-DUF760-1* or *g-DUF760-2* in the mutants of *clpc1-1* and *clpr2-1* showed similar virescent and smaller phenotypes as the respective *clpc1-1* and *clpr2-1* lines (Figure 5A and 6A). We then assessed the protein level of DUF760-1,2 in WT, *clpc1-1,* and *clpr2-1*. Immunoblotting analysis showed that DUF760-1 increased 3 to 5-fold in *clpc1-1* and *clpr2-1* in comparison to WT (Figure 5B). DUF760-2 was not visible in WT but accumulated as very clear bands and at very high levels in both *clpc1-1* and *clpr2-1* (Figure 6B). qPCR results showed that the total mRNA levels of *DUF760-1* and *DUF760-2* (endogenous and transgenic) in the *clpc1-1* and *clpr2-1* are comparable to those in the WT (Figure 5C and 6C). Furthermore, RT-PCR showed that mRNA levels of just the transgenic mRNA product were the same level across the three genetic backgrounds (WT, *clpc1-1* and *clpr2-1*) (Figure 5D and 6D). This rules out that higher transcript levels of *DUF760-1* and DUF760-*2* were the cause of increased accumulation of the two DUF760 proteins in *clp* mutants Collectively, these results are consistent with DUF760-1 and DUF760-2 being substrates of the CLP protease system.

**Figure 5.**
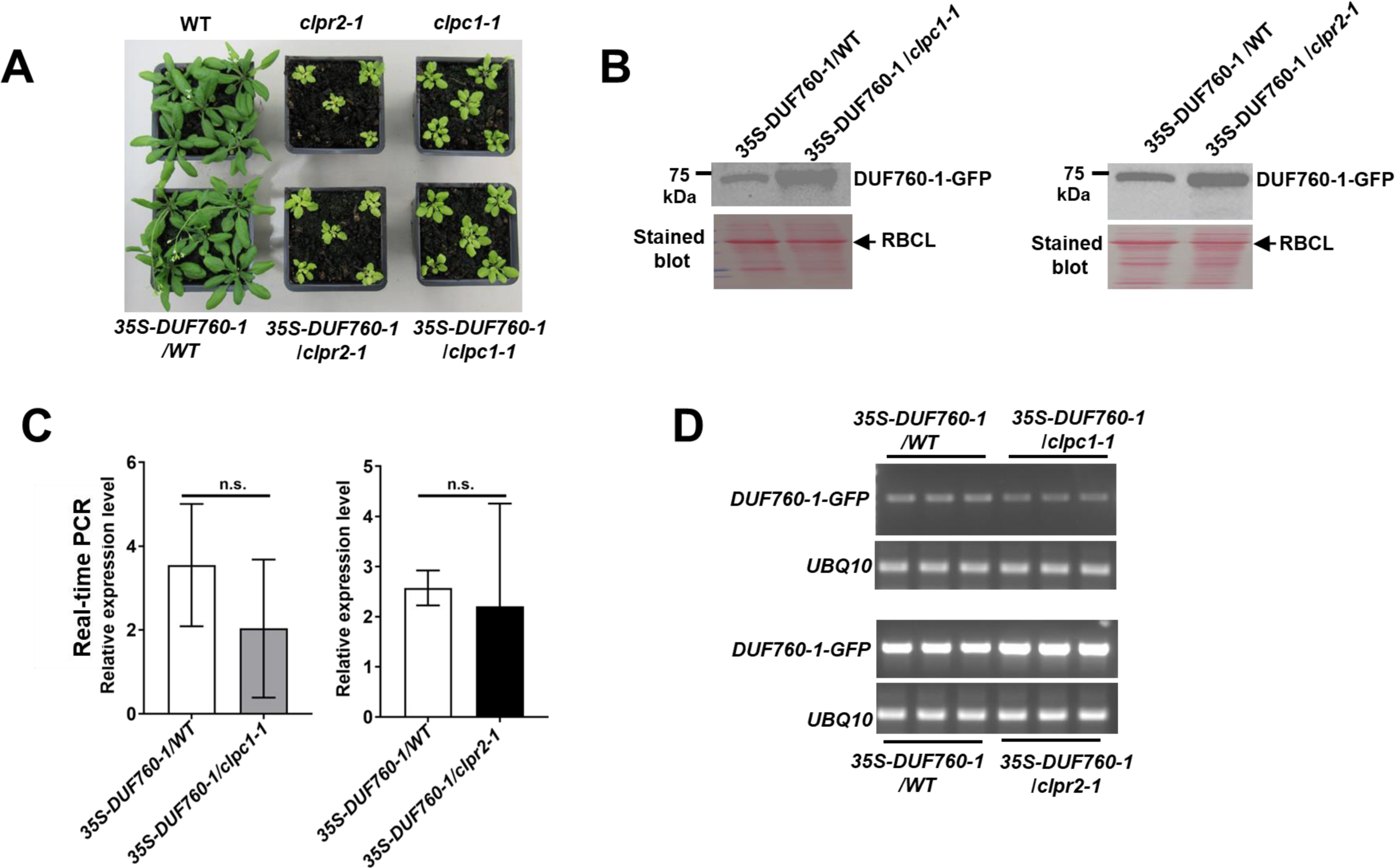
DUF760-1 protein levels in *clpc1-1* and *clpr2-1*. **(A)** Phenotype analysis of *35S-DUF760-1-GFP* in the WT, *clpc1-1,* and *clpr2-1* grown in long day (12h day/12h night, 21°C/19°C) in 100 μmol photons m^−2^s^−1^ of PAR (Photosynthetically Active Radiation). No visible differences were detected. **(B)** Immunoblot analysis of DUF760-1 protein levels. Total protein extracts from 1-week-old seedlings were analyzed by immune blotting using an anti-GFP antibody. **(C)** Real-time PCR of the transcript levels of combined endogenous *DUF760-1 and DUF760-1-GFP* in *35S-DUF760-1/WT*, *35S-DUF760-1*/*clpc1-1* and *35S-DUF760-1*/*clpr2-1*. **(D)** RT-PCR analysis of the transcript levels of transgenic *DUF760-1-GFP*. Transcripts were extracted from the plants of *35S-DUF760-1/WT*, *35S-DUF760-1*/*clpc1-1* and *35S-DUF760-1*/*clpr2-1* and amplified 25 cycles by RT-PCR with gene-specific primers and analyzed on an agarose gel. *UBQ10* mRNA was used as an internal control.

**Figure 6.**
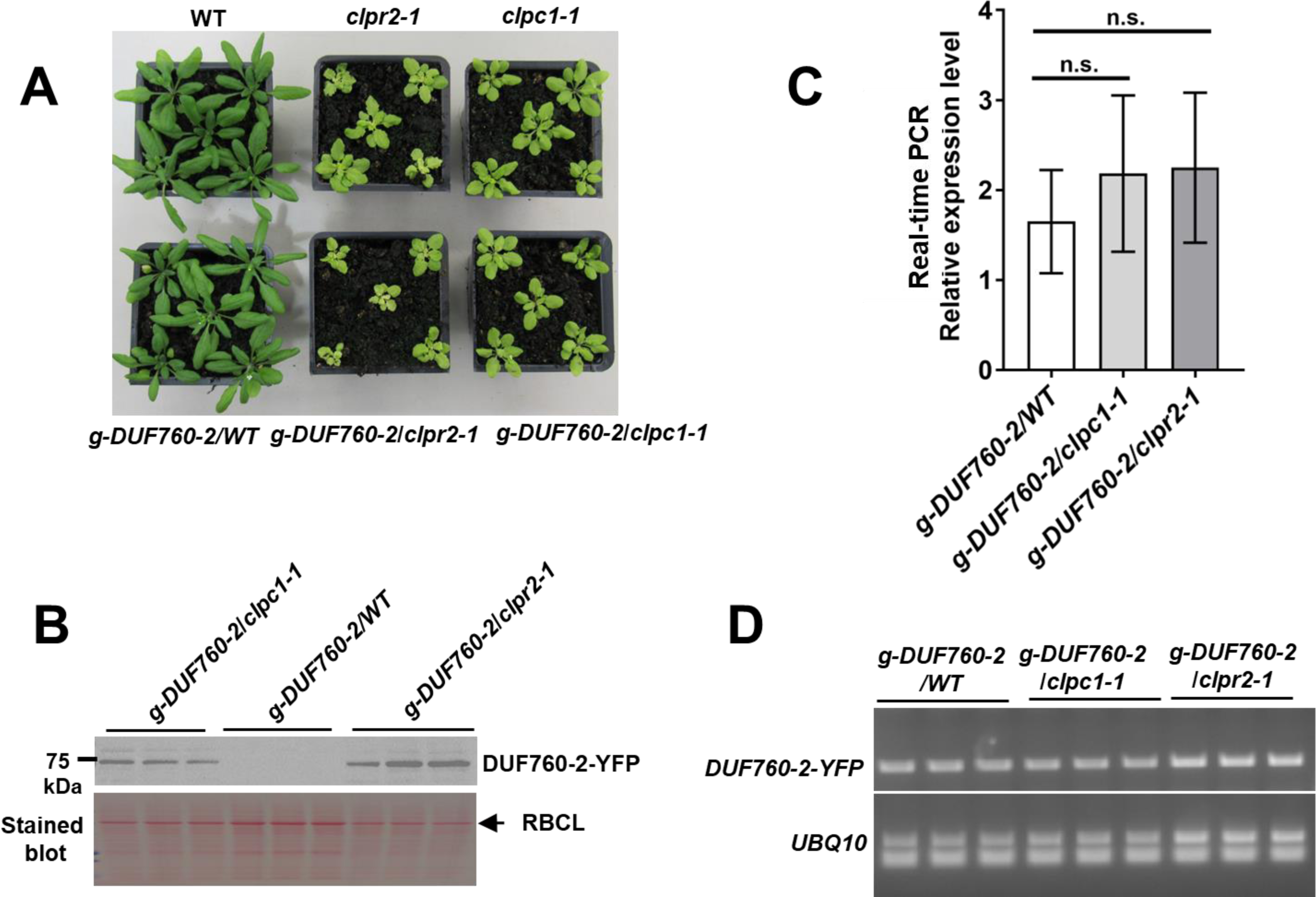
DUF760-2 protein levels in *clpc1-1* and *clpr2-1*. **(A)** Phenotypic analyses of *g-DUF760-2-YFP* in the WT, *clpc1-1,* and *clpr2-1* grown in long day (12h light /12h dark; 21°C/19°C) in 100 μmol photons m^−2^s^−1^ of PAR (Photosynthetically Active Radiation). No visible differences were detected. **(B)** Immunoblot analysis of DUF760-2-YFP protein levels. Total protein extracts from 1-week-old seedlings were analyzed by immune blotting using an anti-GFP antibody. **(C)** Real-time PCR analysis of the transcript levels of combined endogenous DUF760-2 and DUF760-2-YFP in *g-DUF760-2*/WT, *g-DUF760-2*/*clpc1-1*, and *g-DUF760-2*/*clpr2-1*. **(D)** RT-PCR analysis of the transgenic *DUF760-2-YFP* mRNA. Transcripts were extracted from the plants of *g-DUF760-2*/WT, *g-DUF760-2*/*clpc1-1*, and *g-DUF760-2*/*clpr2-1* and amplified 25 cycles by RT-PCR with gene-specific primers and analyzed on an agarose gel. *UBQ10* mRNA was used as an internal control.

### Protein half-life of DUF760-1 and DUF760-2

To determine DUF760-1,2 protein half-lives and their dependence on CLP activity, we performed a CHX chase protein analysis in WT and *clpr2-1* (Figure 7A, B). Figure 7A shows a representative immunoblot with three time points (0, 6 and 12 hrs) of DUF760-1 levels in presence or absence of CHX in WT or *clpr2-1*. DUF760-1 levels were always higher in *clpr2-1* than in WT. In the WT background in presence of CHX, DUF760-1 levels dropped to below 50% after 6 hours and then gradually decreased further over the next 6 hours. In contrast, DUF760-1 levels stayed relatively constant in WT in absence of CHX. In contrast, DUF760-1 levels in *clpr2-1* background remained relatively constants over 12 hours irrespective of the presence of CHX. Figure 7B shows that DUF760-2 accumulated only at very low or undetectable levels in WT at any of the three time points, irrespective of the presence of CHX. In contrast, DUF760-2 was hugely enriched in *clpr2-1* (as also observed in Figure 6B) and barely changed after 3h and 6h during CHX or mock treatments (Figure 7B). These results show that the CLP protease system is required for the degradation of DUF760-1 and 2 and that in particular DUF760-2 has a very high turnover rate due to rapid degradation by the CLP system.

**Figure 7.**
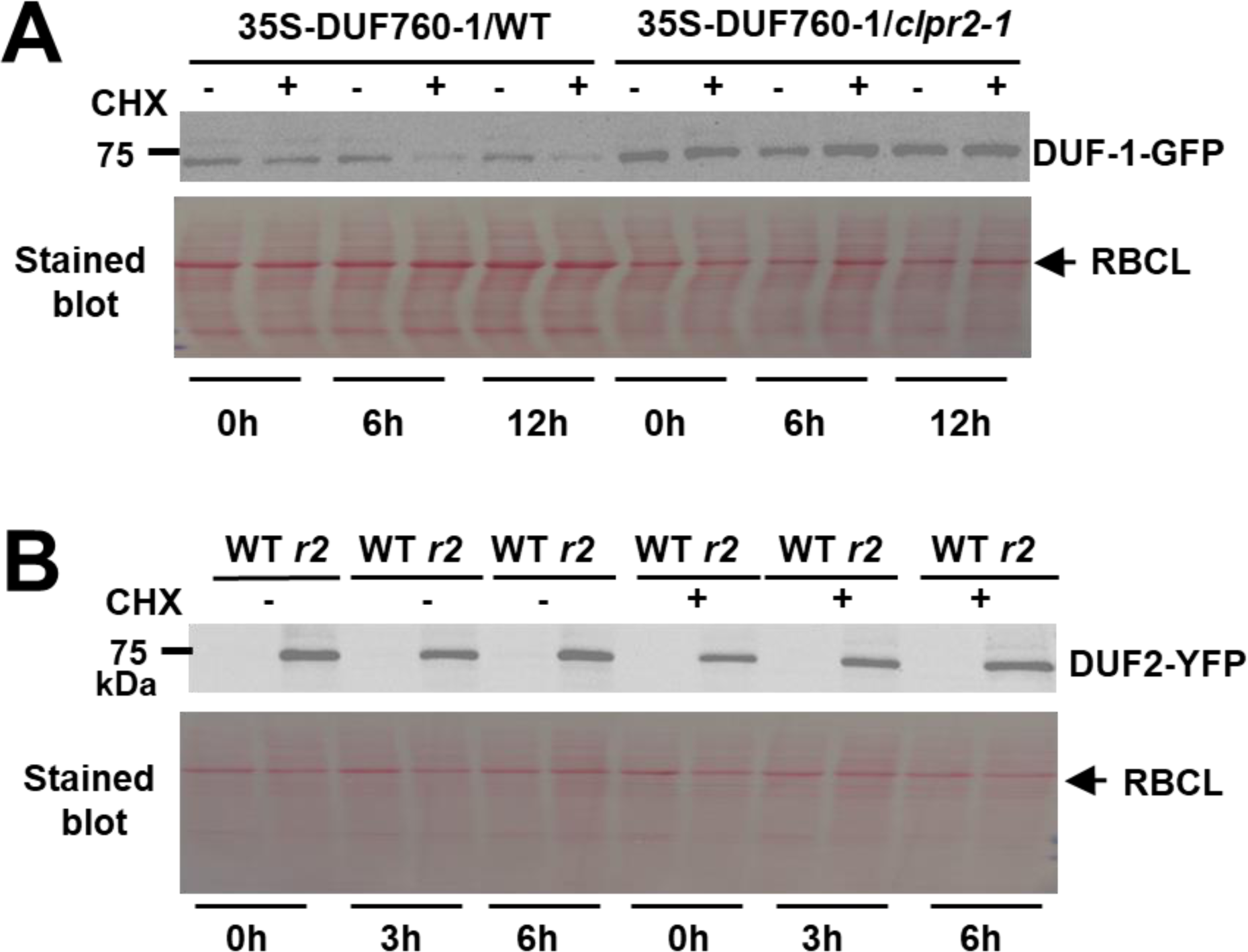
Protein accumulation and half-life of DUF760-1 and DUF760-2 in the WT and the *clpr2-1* mutant. **(A)** Immunoblot analysis of DUF760-1-GFP fusion protein. *35S-DUF760-1-GFP* was expressed in WT and *clpr2-1* mutant. DUF760-1-GFP protein stability was monitored by immunoblotting using an anti-GFP antibody prior to (0 h) and at 6, and 12h following treatment with 300 µM CHX to inhibit cytoplasmic protein translation. Ponceau S staining shows protein loading. **(B)** Immunoblot analysis of DUF760-2-YFP fusion protein. *g-DUF760-2-YFP* was expressed in WT and *clpr2-1* mutant. DUF760-2-YFP protein stability was monitored by immunoblotting using an anti-GFP antibody prior to (0 h) and at 3, and 6h following treatment with 300 µM CHX to inhibit cytoplasmic protein translation. Ponceau S staining shows protein loading.

### DUF760-1 and DUF760-2 directly physically interact with CLPC1

To determine if the two DUF760 proteins directly interact with CLPC1, we performed yeast two hybrid (Y2H) assays (Figure 8). We tested for interaction with the CLPC1 N-terminal portion (aa 39-252; the first 38 aa residues are the cleavable cTP) and the middle portion of CLPC1 (aa 253-635) (Figure 8A). The N-terminal portion includes the N-domain which in CLP chaperones generally serves as a binding site for adaptor proteins and substrates (Erbse et al., 2003; Hanson and Whiteheart, 2005; Lupas and Martin, 2002; Olivares et al., 2018). The CLPC1 middle portion includes the first ATPase domain (Walker B domain) for substrate unfolding and translocation to the core and the uvrB/C motif of unknown function (Figure 8A) (Nishimura and van Wijk, 2015). The Y2H results show that CLPC1-N-Domain but not CLPC1 (middle) in yeast grown on selective medium can mate with DUF760-1 or DUF760-2, demonstrating that the CLPC1-N-Domain is required for the interaction with DUF760-1 and 2 (Figure 8B).

**Figure 8.**
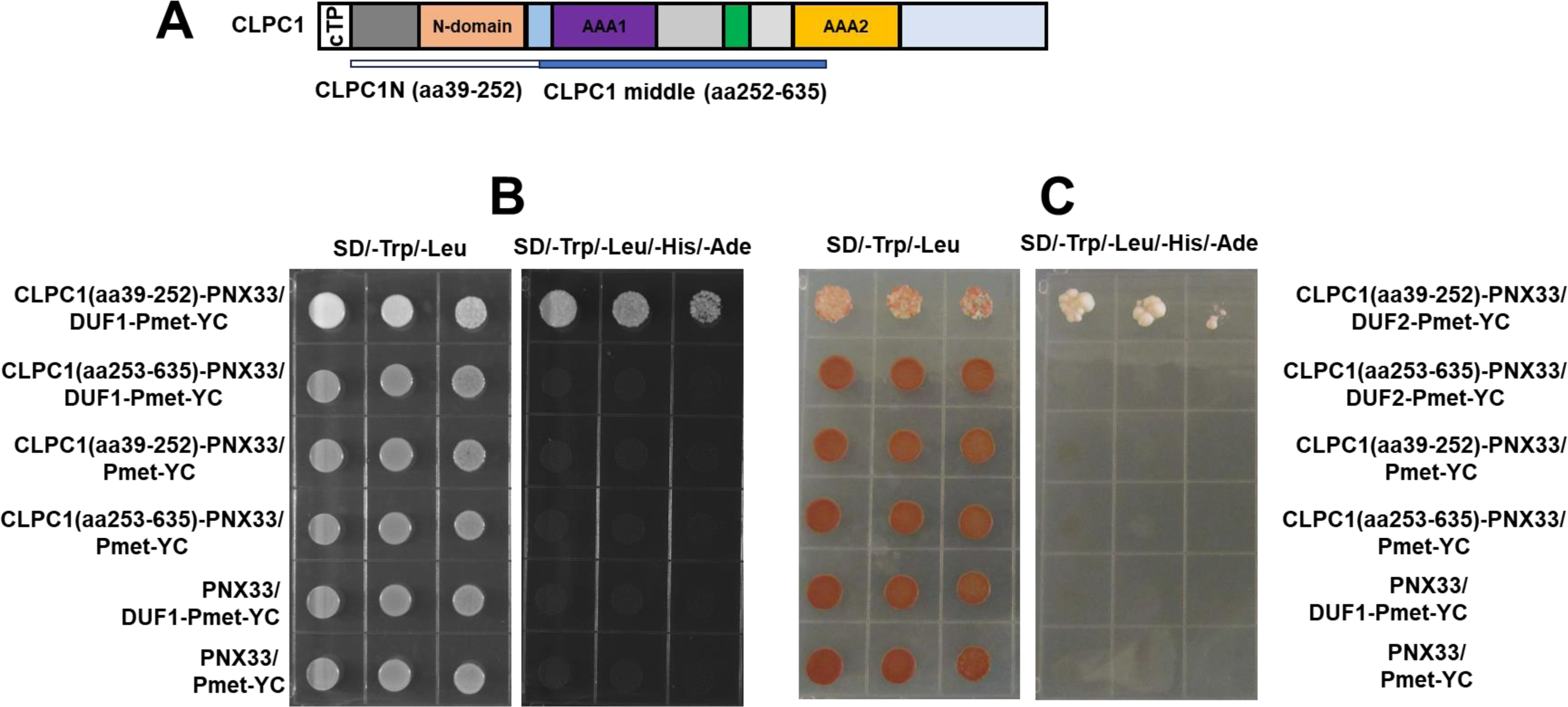
Interaction of DUF760-1 and CLPC1 determined by yeast-2-hybrid assays. **(A)** Schematic structure of CLPC1. CLPC1 protein consists of the chloroplast transit peptide (cTP), the N-domain (with repeats 1 and 2) involved in binding of adaptors (*e.g.* CLPS1) and substrates, the uvrB/C domain of unknown function, and two AAA+ domains (AAA1 and AAA2) involved in ATP-dependent unfolding of substrates. **(B, C)** Yeast two-hybrid assays showing DUF760-1 **(B)** and DUF760-2 **(C)** associate with CLPC1 and the N-domain of CLPC1 is required for their interaction. Interactions were detected by co-expressing pairs of proteins fused to either N-terminal or C-terminal ubiquitin moiety in yeast and spotting onto either nonselective (−LW) or fully selective medium plates (−LWAH) in a series of 10-fold dilutions. Empty vectors expressing Nub and Cub only were used as negative controls, respectively.

To further examine the *in vivo* protein interactions between CLPC1 or CLPC1-TRAP with DUF760-1 and DUF760-2, we conducted bimolecular fluorescence complementation (BiFC) assays in *Nicotiana benthamiana* (Figure 9). We prepared fusion constructs encoding the C-terminal fragment of the yellow fluorescent protein (cYFP) fused to full-length mature *DUF760-1* (DUF760-1-cYFP), *DUF760-2* (DUF760-2-cYFP) or mature *CGEP-S781R* (CGEP-S781R-cYFP) under the control of the 35S promoter. CGEP-S781R is a catalytically inactive chloroplast glutamyl aminopeptidase that operates independently of the CLP chaperone-protease complex (Bhuiyan et al., 2020) and thus serves as a negative control for CLPC1 interactions. Additionally, we generated fusion constructs encoding the N-terminal fragment of YFP (nYFP) fused to the full-length *CGEP-S781R* (CGEP-S781R-nYFP), *CLPC1* (CLPC1-nYFP) or *CLPC1-TRAP* (CLPC1-TRAP-nYFP). When *DUF760-1-cYFP* or *DUF760-2-cYFP* were transiently co-expressed with *CLPC1-nYFP* or *CLPC1-TRAP-nYFP*, we detected strong YFP fluorescence in the chloroplasts, as evidenced by overlap with chloroplast autofluorescence (Figure 9A, B). No fluorescence was detected in the negative controls in which CGEP-S781R was co-expressed with DUF760-1, DUF760-2, *CLPC1* or *CLPC1-TRAP* (Figure 9A, B). Thus, the *in vivo* BiFC results support the direct physical interactions between DUF760-1,2 and CLPC1 observed by *in vitro* Y2H.

**Figure 9.**
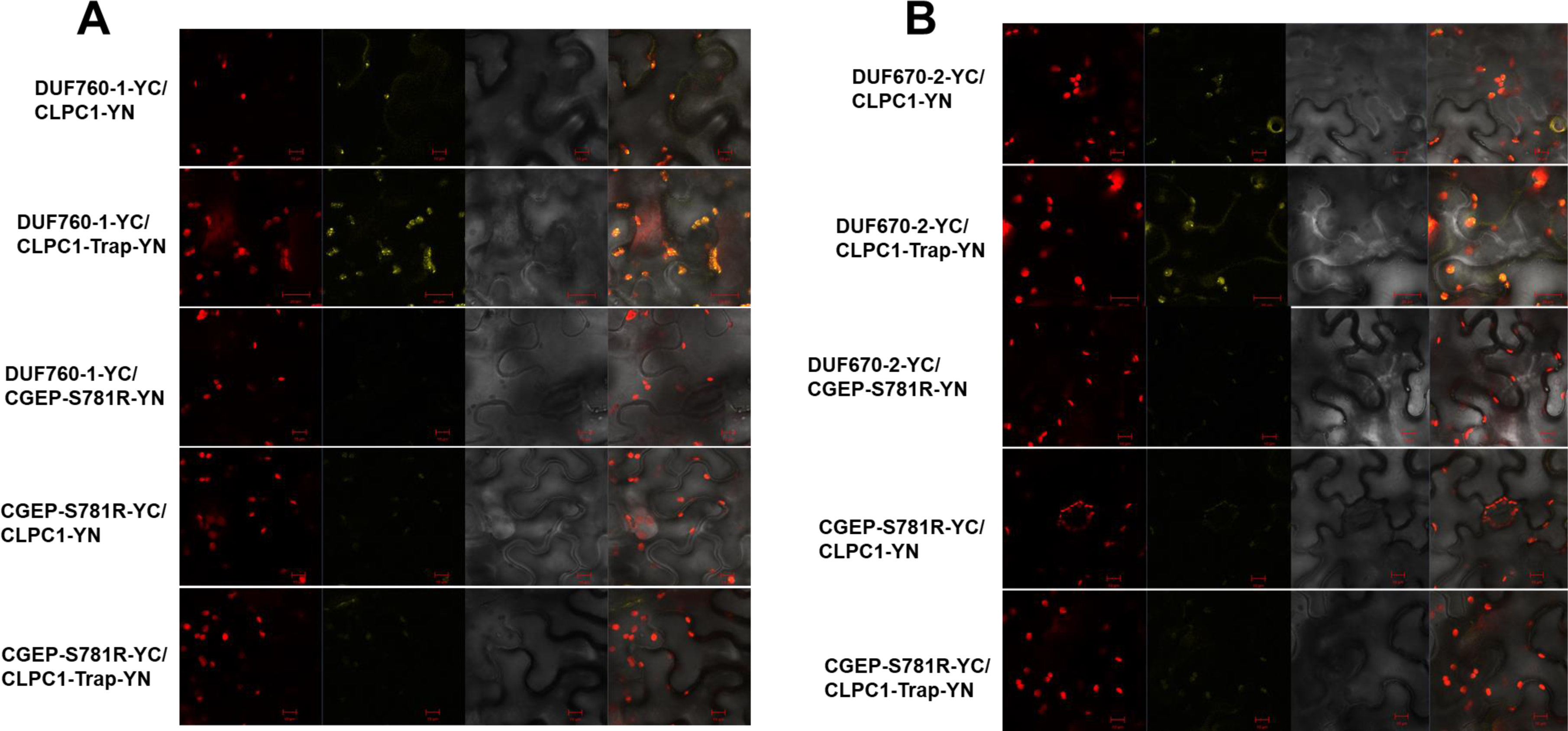
Interaction of DUF760-1 or DUF760-2 with CLPC1 *in planta* by BiFC assays. **(A, B)** DUF760-1 **(A)** and DUF760-2 **(B)** interact with CLPC1-WT and CLPC1-TRAP in chloroplast. Visualization of protein-protein interactions *in planta* by the BiFC assay. Venus confocal microscopy images show *N. benthamiana* leaf epidermal cells transiently expressing constructs encoding the fusion proteins indicated. Bars = 10/20 μm.

### Searching for potential protein interactors to DUF760-1 and DUF760-2 by MSMS in *CLPC1-TRAP* or c*lpc1-1* backgrounds

To identify interactors to DUF760-1 and DUF760-2, we carried out affinity purifications using transgenic 35S-DUF760-1-GFP or 35S-DUF760-2-GFP expressed in four different genetic backgrounds: *CLPC1-WT* and *CLPC1-TRAP* or WT and *clpc1-1.* A transgenic line expressing a chloroplast targeted copy of RECA-GFP in WT background was used as a negative control for unspecific interactions. Affinity eluates were run out on SDS-PAGE gels (Supplemental Figures 7-9 and each lane was cut into 4 or 5 continuous gel slices followed by in-gel digestion with trypsin and recovery of peptides for MSMS analysis and protein identification and quantification based on adjusted spectral counts (adjSPC). We observed increased yield of the affinity enriched DUF760 fusion proteins in the *CLPC1-TRAP* or *clpc1-1* proteins backgrounds as expected and also seen in most of the stained gel images, but except for CLPC1-TRAP, no obvious interactors were identified (data not shown).

## DISCUSSION

The chloroplast CLP chaperone-protease is a soluble and abundant stromal proteolysis machinery that likely acts to degrade hundreds of stromal proteins and likely also proteins degradation products produced by thylakoid or envelope proteases. Therefore the CLP system plays a central role in chloroplast proteostasis, as also demonstrated by embryo or seedling lethal phenotypes when CLP capacity is severely reduced (Bouchnak and van Wijk, 2021; Llamas and Pulido, 2022; Nishimura *et al*., 2016; Nishimura and van Wijk, 2015; Rodriguez-Concepcion *et al*., 2019). The biggest challenge in understanding this central role of CLP is obtaining direct evidence for its substrates and understanding substrate recognition mechanism, *i.e.* what triggers a protein to become a CLP substrate. Previous comparative proteomics studies of a variety of Arabidopsis chloroplast CLP mutants identified numerous proteins that over-accumulated either due to indirect effects (in particular upregulation of the chloroplast chaperones such as HSP70, CPN60, HDP90 CLPB3) or because they are CLP substrates, and their degradation rates is much reduced in these CLP mutants (Kim et al., 2015; Kim et al., 2013; Nishimura *et al*., 2013; Wu et al., 2013; Zhang et al., 2018; Zybailov et al., 2009). *In vivo* trapping with transgenic modified CLPC1 chaperones has allowed capturing potential substrates (Montandon *et al*., 2019; Rei Liao *et al*., 2022) including the DUF760-1,2 proteins which are the focus of this study. Our objective was to determine if these *in vivo* trapped proteins are indeed CLP substrates. Here we have shown that DUF760-1,2 are indeed substrates, thus also demonstrating that the *in vivo* CLPC1 trapping approach is an effective way to discover new substrates, including proteins with short half-lives.

We provided several lines of evidence that both DUF760-1,2 proteins accumulate in green plant tissue types which are consistent with mRNA-based expression patterns. Surprisingly, whereas mRNA levels of DUF760-1,2 are very similar (based on large scale public data collected in ePlant at https://bar.utoronto.ca/eplant/), the protein levels of DUF760-2 are much lower than of DUF760-1, as illustrated by confocal microscopy as well as immunoblotting. This study resolved this mismatch between mRNA and protein level by demonstrating that DUF760-2 has a very short half-life (minutes) compared to the much longer half-life of DUF760-1 (4-6 hours). However, we also presented several independent lines of evidence that both DUF760 proteins are degraded by the CLP system, and both can directly interact with the N-domain of CLPC1. This N-domain has a conserved function in substrate binding across CLP chaperones in eukaryotes and prokaryotes (Erbse *et al*., 2003; Hanson and Whiteheart, 2005; Lupas and Martin, 2002; Olivares *et al*., 2018).

Both DUF760-1 and DUF760-2 have so-called DUF760 domains – the function of this domain is unknown and in fact this DUF is also rather poorly defined. Evaluation of predicted DUF760 structures from alpha-fold (https://alphafold.ebi.ac.uk/) did not provide any clues for molecular functions (not shown). Furthermore, despite both DUF760 proteins having these domains of unknown function, there is no evidence that the DUF760 proteins have similar functions. Indeed, mRNA-based co-expression data show that DUF760-1,2 have very few overlapping co-expressors, even if most co-expressors encode for proteins targeted to plastids (Supplemental Table 1). Indeed, most genes encoding for plastid proteins also co-express with genes encoding for plastid proteins (Majsec *et al*., 2017).

This study provides several independent lines of evidence that the DUF760-2 protein has a very short half-life and that this high turnover rate is strictly dependent on the CLP system. First, DUF760-2 was identified as a highly enriched protein in the CLPC1-TRAP affinity eluates (Rei Liao *et al*., 2022) whereas the protein is rarely detected and only with low numbers of match MSMS spectra across many independent proteomics datasets, as reported in Arabidopsis PeptideAtlas (https://peptideatlas.org/builds/arabidopsis/). Second, the DUF760-2 protein was very hard to detect by immunoblotting in WT Arabidopsis plants, but very easy to detect at high levels in both *clpr2-1* and *clpc1-1* mutants. Confocal microscopy in transgenic lines of *g-DUF760-1-YFP* or *g-DU760-2-YFP* supported the immunoblotting results and also provides further evidence that both proteins accumulate in chloroplasts. Interestingly, DUF760-2 was also detected at low level in non-photosynthetic plastids in root tips both by immunoblotting and confocal microscopy, whereas DUF760-1 was not detected in roots.

The very short-half-life of DUF760-2 (minute scale) as compared to the 4-6 hours half-life of DUF760-1 is intriguing. The PPR protein GUN1 (Richter et al., 2023; Shimizu and Masuda, 2021; Tadini et al., 2020; Wu and Bock, 2021) is another example of a chloroplast protein with a demonstrated short half-life (∼4 hours), and interestingly is also dependent of CLPC1 for its degradation (Wu et al., 2019). However, despite tremendous efforts by a number of laboratories there is still no consensus on the direct function of GUN1 (as also reflected in the numerous reviews cited above). Interestingly it was recently suggested that the degron in Arabidopsis GUN1 lies in its N-terminus of the mature protein, because GUN1 in tomato does not have a short-half life and it lacks the N-terminal region found in Arabidopsis GUN1, although it can completely complement the Arabidopsis *GUN1* mutants (Su et al., 2024). An important question is how DUF760-1 and DUF760-2 (and GUN1) are recognized as a substrate by CLPC1; in other words what are the degrons of these proteins and are these degrons conditional?

Other plant proteins with a demonstrated very short half-life (minute scale) are members of the ERFVII transcription factor family involved in oxygen sensing (Barreto et al., 2022; Gibbs et al., 2015; Hammarlund et al., 2020; van Dongen and Licausi, 2015; Weits et al., 2021). These ERFVII proteins have a short half-life under normal oxygen concentration (normoxia) because they are degraded by the cytosolic or nuclear proteasome through the N-degron pathway. At low cellular oxygen concentrations these transcription factors stabilize because the enzymatic oxidation is slowed down (Abbas et al., 2022). These ERFVII proteins have a cysteine in the 2^nd^ position from the N-terminus. After removal of the start methionine by methionine amino peptidases, these N-terminal cysteines are enzymatically oxidized by Plant Cysteine Oxidases (PCDs) which is then followed by enzymatic arginylation (Hammarlund *et al*., 2020; White et al., 2017). The arginylated N-terminus is then recognized by specific E3 ligases, resulting in polyubiquitination and degradation by the proteasome. However, these ERVII proteins, as well as the PCDs, arginylase, E3 ligases and proteoasome are all localized outside of chloroplasts and hence this N-degron pathway is unlikely to contribute to the short half-life of DUF760-2.

Future studies will be focused on identification of the DUF760-1 and DUF760-2 degrons, including the role of possible adaptors that can enhance delivery to the CLPC chaperone. Finally, we note that our prior CLP1 trapping studies identified a number of other highly enriched proteins, some of which normally accumulate at very low steady state levels (Montandon *et al*., 2019; Rei Liao *et al*., 2022). The current study shows that these enriched proteins provide a great resource to explore other candidate CLP substrates and possibly identify novel degrons.

## METHODS

### T-DNA insertion lines

*Arabidopsis thaliana* ecotype *col-0* was used in this study. The *duf760-1-1* (SALK_030328), *duf760-1-2* (SALK_066581), and *duf760-2-1* (SALK_124643) mutants were obtained from the Arabidopsis Biological Resource Center (ABRC). The *clpc1-1* and *clpr2-1* mutants and the 35S-cpRECA/WT lines were described previously (Kohler et al., 1997; Nishimura *et al*., 2013; Rudella et al., 2006). The positions of the T-DNA insertions were confirmed by DNA sequencing. All plants were grown at 22°C under constant (24 h) light or under 16/8-h light/dark cycle (∼100 µmol.m^−2^.s^−1^ of fluorescent lamp) and 60% humidity conditions unless stated otherwise.

### Growth on antibiotics of T-DNA insertion lines

For antibiotic treatments, *duf760-1-1*, *duf760-1-2*, *duf760-2-1*, and the WT were grown with or without antibiotic (CAP and CHX) in half-strength Murashige and Skoog plates (0.5× Murashige and Skoog salts [Sigma-Aldrich], 0.6% [w/v] Phytoblend [Caission Laboratories], and 2% [w/v] Suc, pH 5.7) under 10-h-light (∼100 µmol.m^−2^.s^−1^ of fluorescent lamp) at 22°C / 14-h-dark at 22°C cycles for 14 or 20 to 21 days.

### Construction of transformation vectors and generation of transgenic plants

For protein expression pattern experiments of DUF760-1 and 2, the genomic DNAs of *DUF760-1* and *2* (without the stop codon) were cloned with specific primers (Supplemental Table 2) into the Gateway compatible vector pCR™8/GW/TOPO (Invitrogen) and then transferred to the vector pGWB540 with the YFP at the C-terminus using LR clonase (Invitrogen). For transgenic *Arabidopsis* plants overexpressing *35S-DUF760-1-GFP* or *35S-DUF760-2-GFP* in the wild type, the CDS of *DUF760-1* or 2 were subcloned in the gateway entry vector pCR™8/GW/TOPO and finally inserted by LR reaction in the plant expression binary PGWB5 vector with the GFP at the C-terminus, which carries a 35S promoter and a hygromycin B resistance gene for the selection of the transformants, yielding the vector *35S-DUF760-1-GFP* or *35S-DUF760-2-GFP*. All plasmids generated above were transformed in Agrobacterium tumefaciens GV3101 strain by electroporation following the electroporation device’s manufacturer’s instructions (Bio-Rad). Information on all transgenic lines used in the study is summarized in Supplemental Table 2.

### RNA extraction and qRT–PCR assays

Total RNA was isolated with a RNeasy plant mini kit (Qiagen). To eliminate contaminating genomic DNA, RNA was treated with DNase (TURBO DNA-free Kit; Ambion). The first strand was synthesized from equal amounts of total RNA with Superscript III Reverse Transcriptase (Invitrogen). qRT-PCR analyses were performed as described previously (Jiang et al., 2023). The thermal cycling conditions were as follows: 42 cycles of 95 °C for 10 s, 56 °C for 15 s, and 72 °C for 20 s. Specific primers for each gene are listed in Supplemental Table 2. Results were normalized using *UBQ10* as the endogenous control. Three biological replicates were analyzed for each experiment. Transcript levels of *DUF760-1,2* in the t-DNA lines were determined using reverse transcriptase PCR (RT-PCR). The reaction mixture (10µl) contained the template (0.5–2.5 µg), 0.2 µM of each primer, and Promega GoTaq™ G2 Green Master Mix. The amplification conditions were as follows: 95 °C for 4 min, followed by 25 cycles of 95 °C for 30 s, 55 °C for 30 s, and 72 °C for 1 min, and a final extension step at 72 °C for 5 min.

### Bimolecular fluorescence complementation

BiFC assays were carried out as described before (Zhou et al., 2015). Briefly, the various proteins were fused to the N/C-terminal halves of EYFP (nEYFP/ cEYFP) using the pBatTL-B-sYFPc gateway vectors (gift from Joachim Uhrig; MPIPZ Cologne, Germany). The various constructs were introduced in *Agrobacreium tumefaciens* strain GV3101. Overnight cultures were diluted to an OD600 of 0.5 in resuspension buffer (10 mM MgCl2, 10 mM MES (pH 5.7) and 100 μM acetosyringone and injected into 4–6-week-old *N. benthamiana* leaves with a syringe. Forty-eight hours later, YFP fluorescence was detected using a CLSM Zeiss LSM 700 laser scanning confocal microscope with excitation wavelength at 488 nm and emission filter at 520 nm. Chloroplasts were excited with the blue argon laser (488 nm), and emitted light was collected at 680–700 nm, image acquisition and the export of TIFF files.

### Yeast two-hybrid assay

The split ubiquitin system was used as described previously (Zhou et al., 2015). The CDS (without stop codon) of CLPC1 (aa39-252), and CLPC1 (aa253-635) were cloned to make Nub plasmids using LR reactions. The cDNA sequence of *DUF760-1* and *2* were cloned into the Cub-expressing vector. Plasmids were transformed into yeast strain THY.AP4 (Nub) or THY.AP5 (Cub) and mated with each other. Interactions were examined by placing yeast strains with a series of dilutions on selection medium lacking leucine, tryptophan, adenine, and histidine (−LWAH) after 2 days of growth at 29°C.

### Pigment assays

To quantify chlorophyll and carotenoid contents, seedlings were ground to a powder in liquid N2 and extracted with 80% acetone overnight in the dark at 4°C, followed by removal of cell debris by centrifugation for 10 min at ∼15,000g. The chlorophyll and carotenoid quantifications were according to (Lichtenthaler, 1987).

### Protein extraction and immunoblot analysis

For western blots, about 20 mg of tissue from 7-day-old seedlings was ground into fine powder in liquid nitrogen with a TissueLyser system (QIAGEN). Total proteins were extracted using denaturing buffer (100 mM Tris-HCl pH 7.5, 4% SDS (sodium dodecyl sulfate, Sigma-Aldrich), 20% glycerol, 0.1M DTT, and 0.01% bromophenol blue) in a 1:5 tissue: buffer (w/v) ratio by boiling for 10 min at 94 °C. Samples were centrifuged at 15,000g for 5 min at room temperature and the supernatants were electrophoresed on a sodium dodecyl sulfate–polyacrylamide gel electrophoresis (SDS–PAGE) mini-gel to separate the proteins. For Ponceau-S staining, nitrocellulose membranes were incubated in Ponceau-S solution (40% methanol (vol/vol), 15% acetic acid (vol/vol), 0.25% Ponceau-S). The membranes were destained using deionized water. Immunoblots were performed with specific antibodies, and chemiluminescence-based signal detection was done with the ChemiDoc Touch Imaging System (Bio-Rad). Antibodies were obtained from commercial suppliers (GFP from Genescript). Band intensities were determined using the Image J.

### CHX chase assays for half-life analysis

Plant material for CHX chase assays (to determine the half-life of nucleus-encoded proteins) was produced by germinating seeds and growing the resulting seedlings for 7 d in long-day conditions in petri dishes on 0.5× MS medium with netting. Seedlings were then transferred to 12 well culture plates with liquid 0.5× MS medium and incubated under slow agitation. The medium was supplemented with 300 µM CHX in 0.03% DMSO or only 0.03% DMSO for mock controls. Samples were collected at different time points and snap frozen in liquid nitrogen for further analyses.

### Confocal Microscopic Analyses

Confocal laser-scanning microscopy (Zeiss 710 Confocal) was used to determine the subcellular localization of GFP and chlorophyll fluorescence. For GFP fluorescence, the excitation wavelength was 488 nm, and emission was detected with a 495-530 nm filter. For detection of chlorophyll fluorescence, a 650- to 702-nm filter was used.

### Protein-GFP/YFP affinity purification

For GFP pull-down assays, 10 g of fresh plant material were ground in liquid nitrogen to a fine powder and vortexed in 10 ml extraction buffer (EB; 50 mM Hepes-KOH pH 8.0; 15% glycerol, 10 mM MgCl2, 75 mM NaCl, 0.32 mg avidin/ml EB, and 250 μg/ml pefablok serine protease inhibitor). The suspension was filtered through four layers of Miracloth (∼25 μm, Millipore), followed by centrifugation for 1.5 h at 25,000 rpm in a SW28 rotor at 4 °C to remove larger particles. The supernatants were collected and aliquoted and directly used for affinity purification on ChromoTek GFP-Trap® Agarose (Cat No. gta). Extracts were incubated with GFP-trap beads on ice for 1 h during which samples were rotated every 5 min. Beads were then washed three times with 1 ml EB without protease inhibitor. Proteins were eluted by incubation of the beads for 5 min 0.1% TFA (with intermittent vortexing) immediately followed by addition of Tris-HCl buffer (1 M, pH 8.0) to the eluates and beads to raise the pH to 7. Eluates and beads were then stored for analysis by SDS-PAGE, in-gel digestion, and MSMS.

### Protein analysis by MSMS

GFP affinity eluates were separated by SDS-PAGE on Biorad Criterion Tris-HCl precast gels (10.5-14% acrylamide gradient). Each of the SDS-PAGE gel lanes were cut into consecutive gel slices (5-6 per lane), followed by reduction, alkylation, and in-gel digestion with trypsin (Rei Liao *et al*., 2022). The peptides were resuspended in 2% formic acid and analyzed using a QExactive mass spectrometer equipped with a nanospray flex ion source and interfaced with a nanoLC system and autosampler (Dionex Ultimate 3000 Binary RSLCnano system) as described in (Rei Liao *et al*., 2022) with the exception that AGC target values were always set at 1 x 10^6^ for the MS survey scans and maximum scan time 30 ms and 5.10^5^ for MSMS scans and maximum scan time 50 ms. Data processing, search and analyzes was as described in (Rei Liao *et al*., 2022).

## AUTHOR CONTRIBUTIONS

B.Y. carried out most experiments and K.J.V.W. carried out the mass spectrometry analysis; B.Y. and K.J.V.W. developed this project and wrote the paper.

## ACKNOWLEDGEMENTS

We thank Pratyush Routray for advice and discussions, Marissa Annis for evaluation of DUF760 structures and discussions and Giulia Friso for advice on mass spectrometry analysis. We thank Dr. Lili for supplying the plasmids for yeast two hybrid assays. We thank Dr. Joachim Uhrig for supplying the plasmids for bimolecular fluorescence complementation (BiFC) assay.

## SUPPLEMENTAL INFORMATION

**Supplemental Table 1.** Top 50 mRNA co-expressors of DUF760-1 and DUF760-2 obtained from the ATTEDII database (https://atted.jp/) and annotated according to PPDB http://ppdb.tc.cornell.edu/.

**Supplemental Table 2.** Transgenic lines and oligos for genotyping, Q-PCR, and RT-PCR.

**Supplemental Figure 1.**
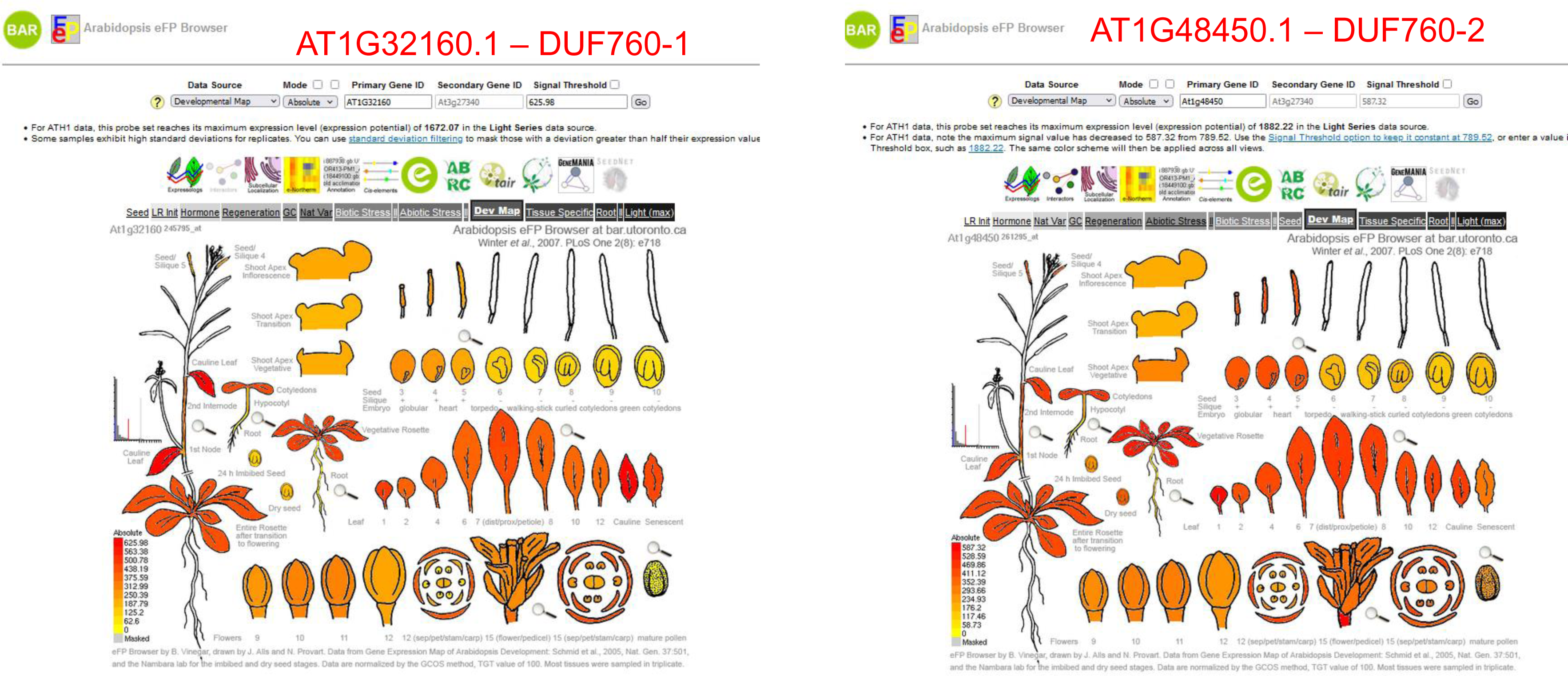
Transcript levels of *DUF760-1* and *2* in different organs at different developmental stages. Relative mRNA accumulation of *DUF760-1* and *2* in different organs at different developmental stages. The images are based on publicly available data at the *Arabidopsis* eFP Browser.

**Supplemental Figure 2.**
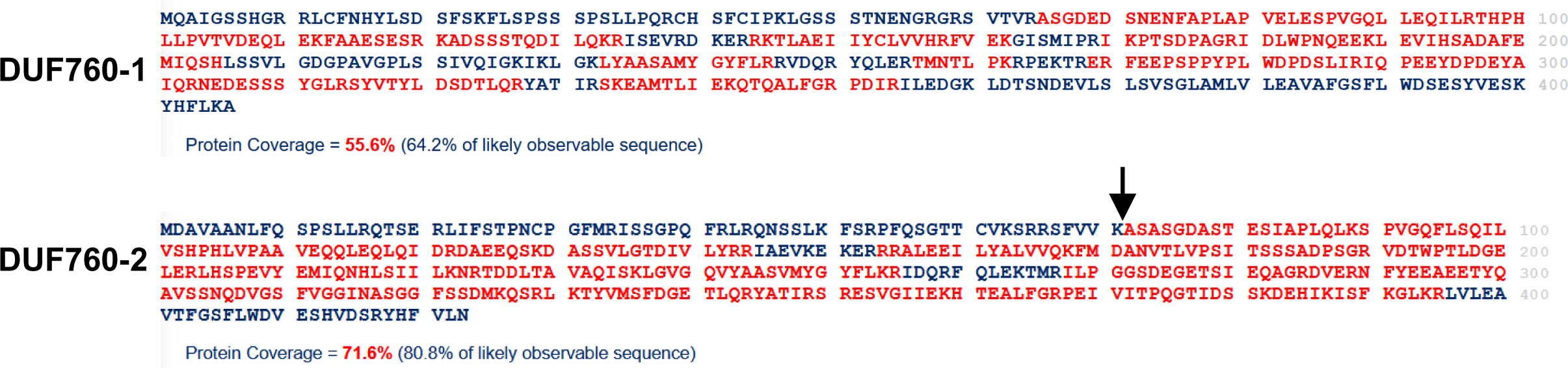
Amino acid sequences of DUF760-1 and 2 and observed residues in red by MSMS as collected in the Arabidopsis PeptideAtlas. The arrow indicates the cleavage site of the cTP as predicted by TargetP.

**Supplemental Figure 3.**
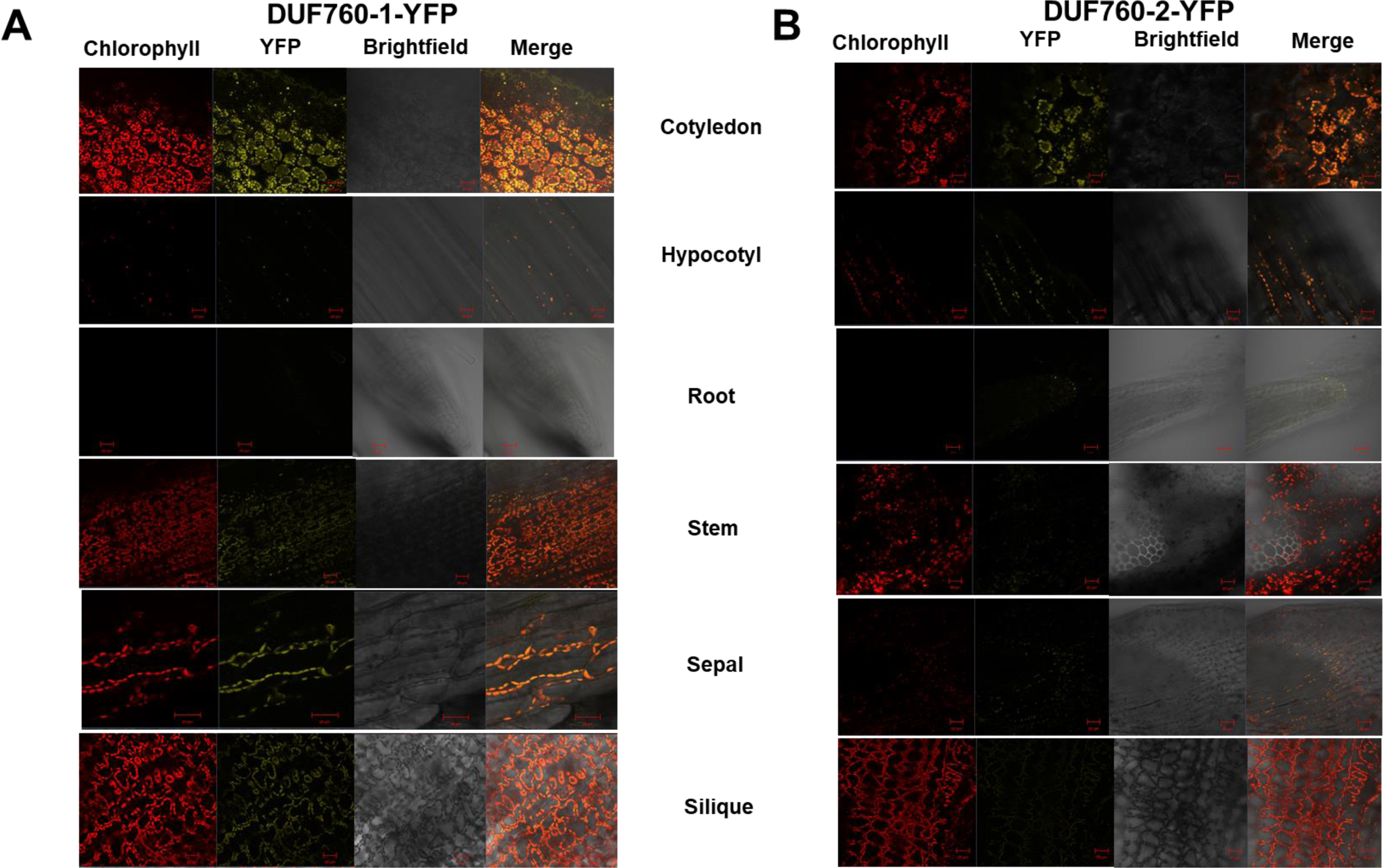
Expression pattern of DUF760-1 and 2 in *Arabidopsis* plants in different tissues. **(A)** Accumulation of DUF760-1-YFP in *Arabidopsis* plants in different organs. YFP signal was detected in the leaves of the transgenic plants expressing *g-DUF760-1-YFP* in the WT in different tissues. **(B)** Accumulation of DUF760-2-YFP in *Arabidopsis* plants in different organs. YFP signal was detected in the leaves of the transgenic plants expressing *g-DUF760-2-YFP* in the WT in different tissues. The *Arabidopsis* plants were grown in chambers in constant light of 100 μmol photons.m^-2^.s^-1^ and a relative humidity of 50%–70%.

**Supplemental Figure 4.**
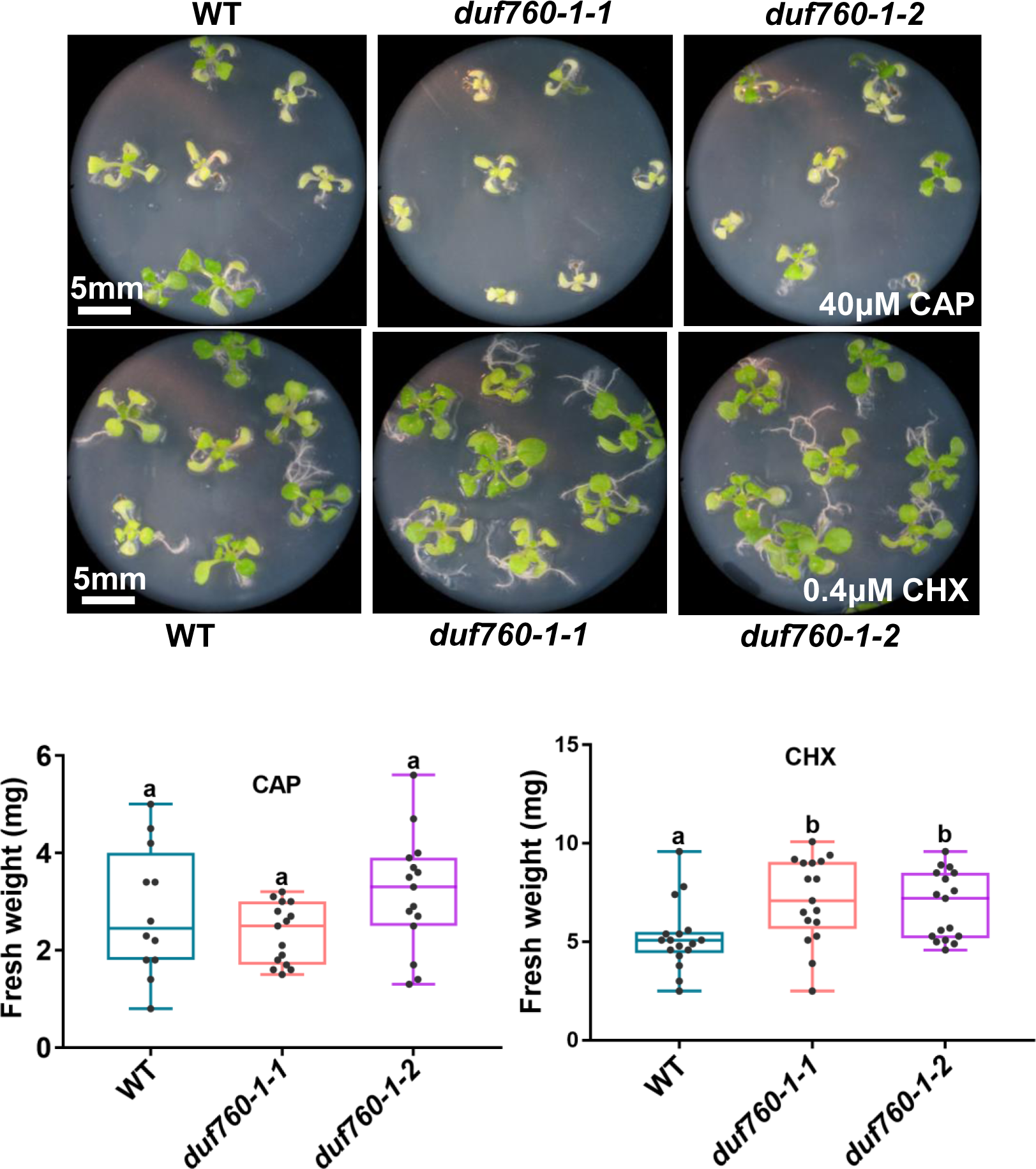
Effects of translation inhibitors on the WT and *duf760-1*. **(A)** Seedling phenotypes of WT, and *duf760-1* lines after treatment with CAP or CHX. Plants were grown for 20 d under short days on half-strength Murashige and Skoog medium containing 1% sucrose and 40μM CAP or 0.4 μM CHX. Bars = 5 mm. **(B)** Effect of CAP or CHX treatment on fresh weight of WT and *duf760-1* seedlings. Error bars indicate SD. Lowercase letters indicate significant differences between multiple groups by one-way ANOVA at p<0.05.

**Supplemental Figure 5.**
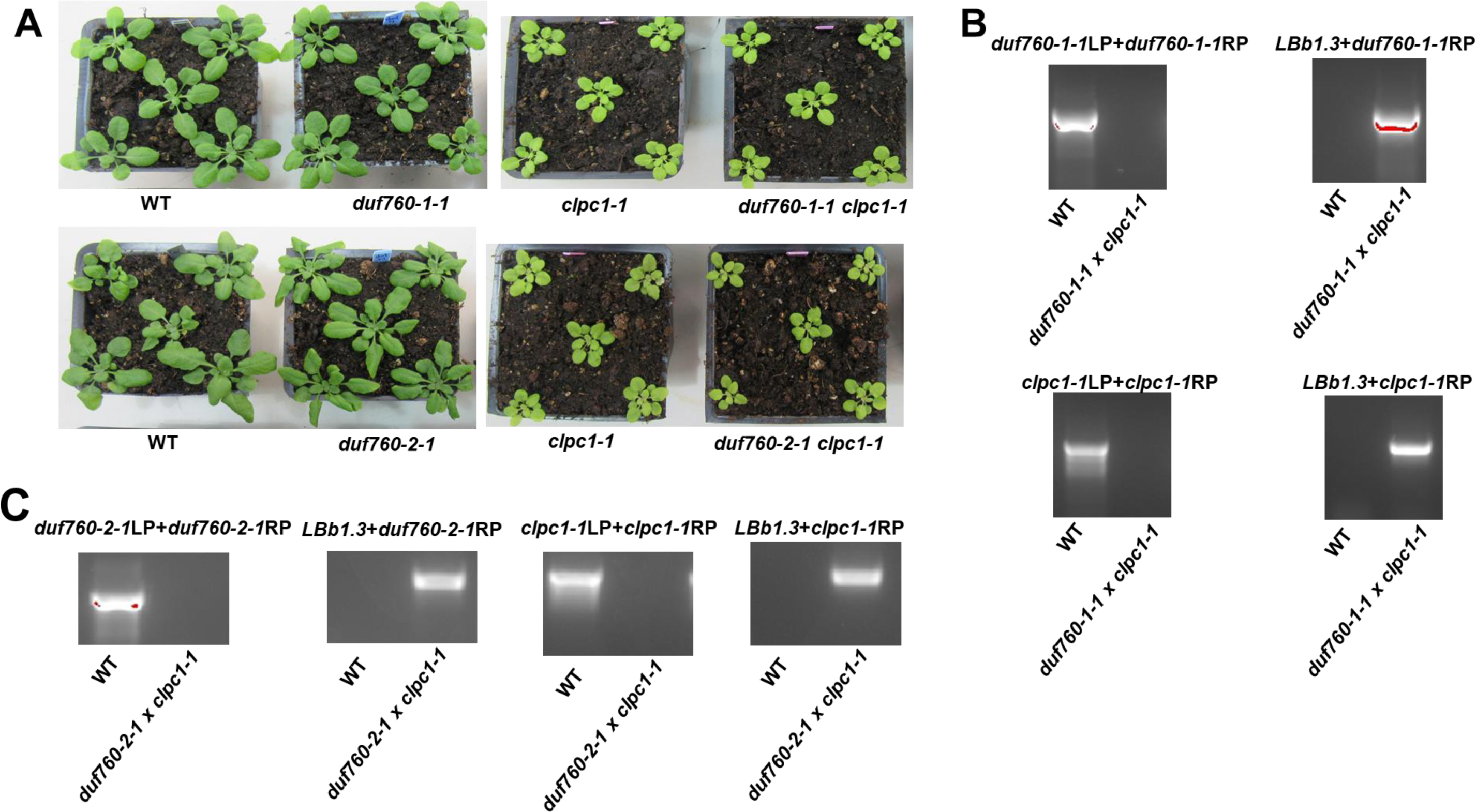
Genetic interaction of *CLPC1* with *DUF760-1* or DUF760-*2*. **(A)** Effect of *duf760-1* or *2* on *clpc1-1*. Homozygous single and double mutant plants were grown in long day (12h day/12h night, 21°C/19°C) in presence of 100μmol m^−2^s^−1^ of PAR (Photosynthetically Active Radiation). **(B, C)** PCR genotyping for analysis of these single and double mutant plants at DNA level.

**Supplemental Figure 6.**
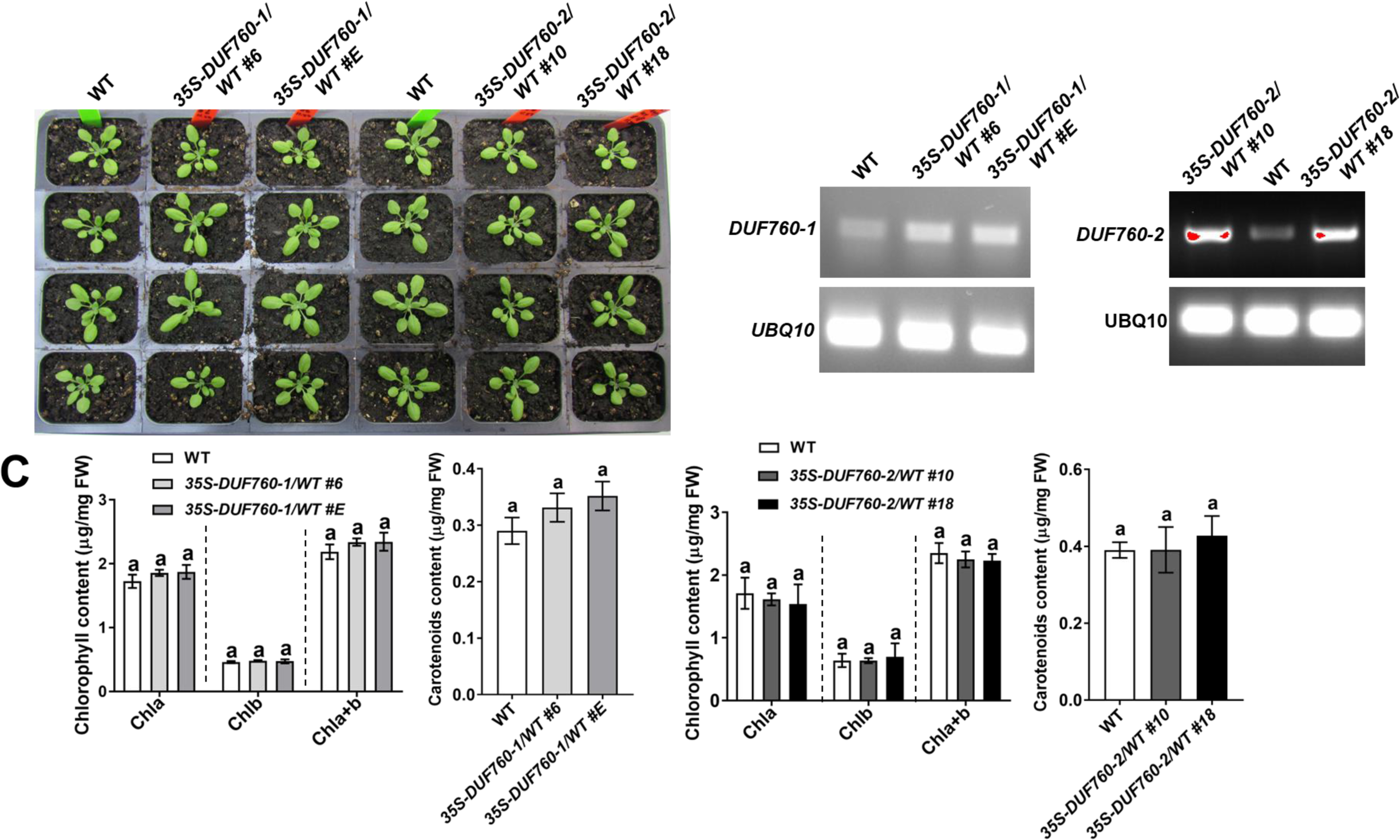
Characterization of the overexpression transgenic lines of *DUF760-1* and *2*. **(A)** Phenotypic analyses of the overexpression transgenic lines of *DUF760-1* and *2*. Plants were grown for 20 d on soil under 16/8-h light/dark cycle at ∼120 µmol photons m^−2^ s^−1^. No visible differences were observed. **(B)** RT-PCR analysis of transgenic lines overexpressing *DUF760-1* and *2* in the WT. Transcripts were extracted from the plants of the WT, and the overexpression transgenic lines of *DUF760-1* and *2* and amplified 25 cycles by RT-PCR with gene-specific primers and analyzed on an agarose gel. *UBQ10* mRNA was used as an internal control. **(C)** Pigment accumulation in the overexpression transgenic lines of *DUF760-1* and *2* at development stage 1.07. Chlorophyll a+b and total xanthophyll and other carotenoid contents were determined on a fresh weight basis. Data are the average of three biological repeats. Error bars indicate SD. lowercase letters indicate significant differences between multiple groups by one-way ANOVA at p<0.05.

**Supplemental Figure 7.**
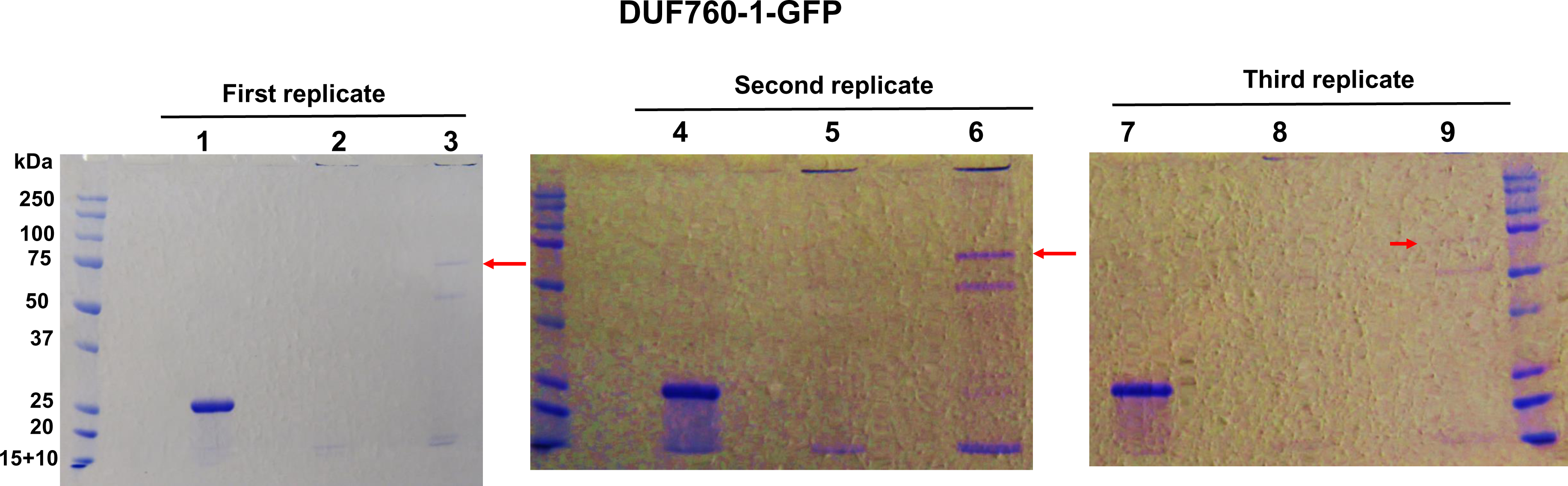
Pull-down experiment of DUF760-1-GFP in *CLPC1-WT* and *CLPC1-Trap.* There are three independent replicates as indicated. Lanes 1,4,7 are *35S-RecA-GFP/*WT, lanes 2,5,8 are *35S-DUF760-1-GFP*/*CLCP1-WT*, and lanes 3,6,9 are *35S-DUF760-1-GFP/CLPC1-TRAP.* Red arrow indicates DUF760-1-GFP fusion protein. Each lane was cut into 5 consecutive gels slices, resulting in 45 samples, that each were analyzed by MSMS.

**Supplemental Figure 8.**
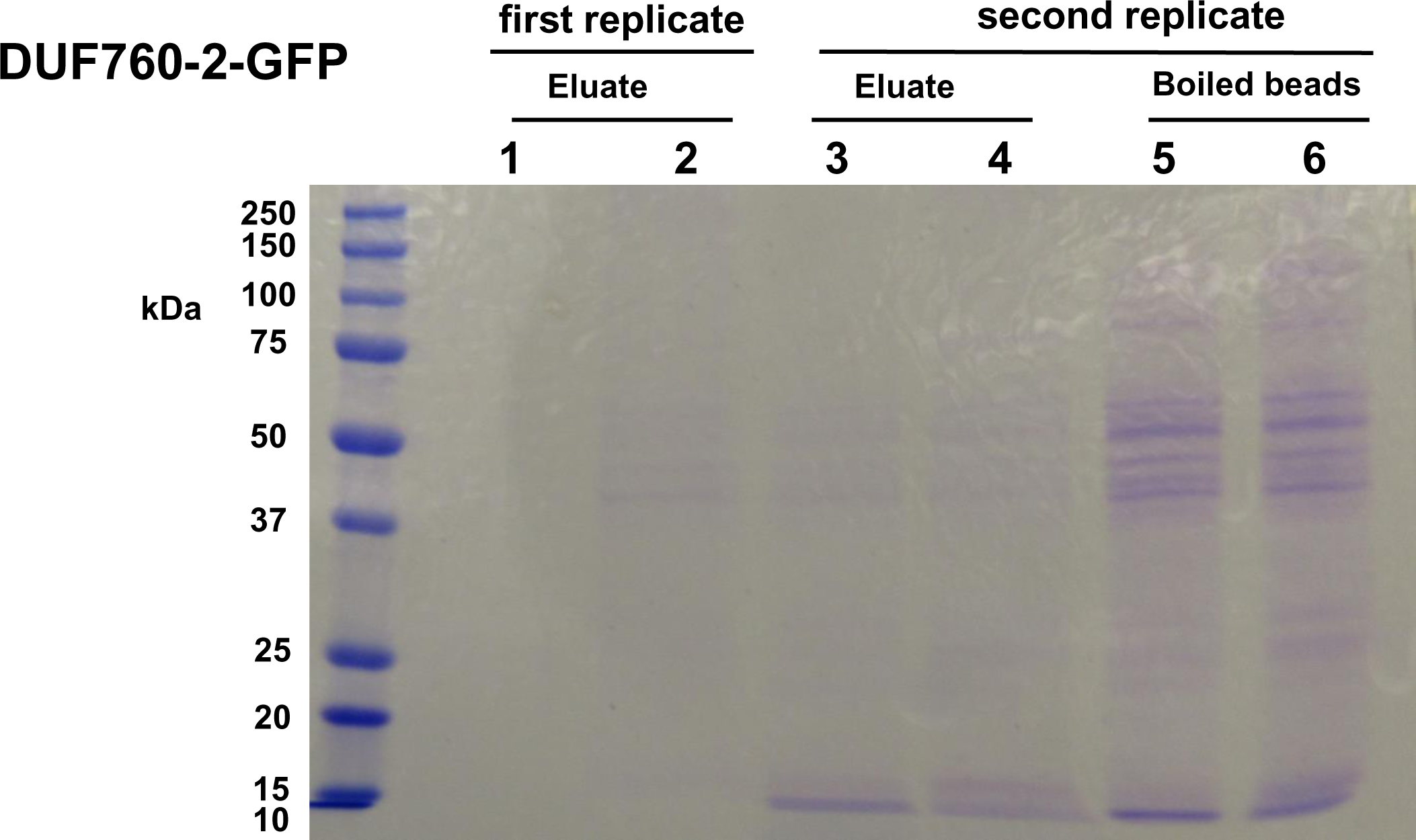
Pull-down experiment of DUF760-2-GFP in *CLPC1-WT* and *CLPC1-TRAP.* Two independent replicates as indicated. Lanes 1,3,5 are *35S-DUF760-2-* GFP/*CLCP1-WT*, and lane 2,4,6 are *35S-DUF760-2-GFP/CLPC1-TRAP.* Each lane was cut into 5 consecutive gels slices, resulting in 30 samples, each of which were analyzed by MSMS.

**Supplemental Figure 9.**
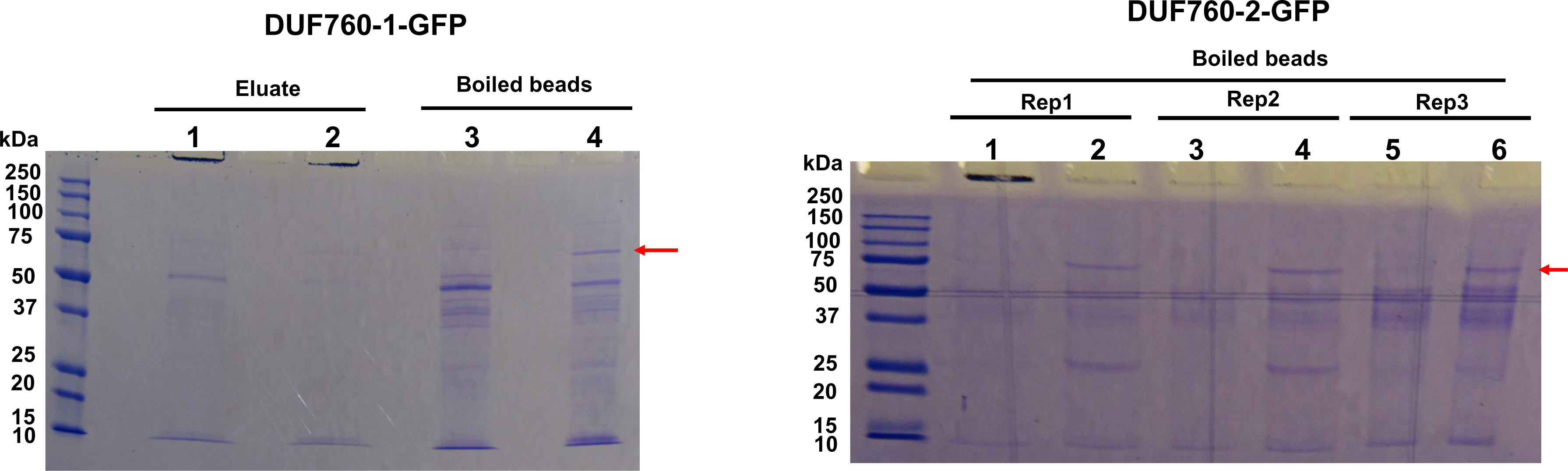
Pull-down experiment of DUF760-1-GFP and DUF760-2-GFP in wild type and *clpc1-1*. **(A)** Lanes 1 and 2 are low pH eluates and lanes 3,4 are proteins released from the low pH treated beads by boiling in SDS. Lanes 1 and 3 are *35S-DUF760-1-GFP*/WT and lanes 2 and 4 are *35S-DUF760-1-GFP*/*clpc1-1*. Red arrow indicates DUF760-1-GFP fusion protein. Each lane was cut into 5 consecutive gels slices, resulting in 20 samples each of which were analyzed by MSMS. **(B)** Three independent replicates as indicated. All samples are from proteins released by boiling the beads in SDS. Lanes 1, 3 and 5 are *35S-DUF760-2-GFP*/WT and lanes 2,4,6 are *35S-DUF760-2-GFP/clpc1-1*. Red arrow indicates DUF760-2-GFP fusion protein. Each lane was cut into 4 consecutive gels slices, resulting in 28 samples each of which were analyzed by MSMS.

## REFERENCES

Abbas, M., Sharma, G., Dambire, C., Marquez, J., Alonso-Blanco, C., Proano, K., and Holdsworth, M.J.(2022). An oxygen-sensing mechanism for angiosperm adaptation to altitude. Nature 606:565–569. 10.1038/s41586-022-04740-y.

Apitz, J., Nishimura, K., Schmied, J., Wolf, A., Hedtke, B., van Wijk, K.J., and Grimm, B.(2016). Posttranslational Control of ALA Synthesis Includes GluTR Degradation by Clp Protease and Stabilization by GluTR-Binding Protein. Plant Physiol 170:2040–2051. 10.1104/pp.15.01945.

Barreto, P., Dambire, C., Sharma, G., Vicente, J., Osborne, R., Yassitepe, J., Gibbs, D.J., Maia, I.G., Holdsworth, M.J., and Arruda, P.(2022). Mitochondrial retrograde signaling through UCP1-mediated inhibition of the plant oxygen-sensing pathway. Curr Biol 32:1403–1411 e1404. 10.1016/j.cub.2022.01.037.

Bhuiyan, N.H., Rowland, E., Friso, G., Ponnala, L., Michel, E.J.S., and van Wijk, K.J.(2020). Autocatalytic Processing and Substrate Specificity of Arabidopsis Chloroplast Glutamyl Peptidase. Plant Physiol 184:110–129. 10.1104/pp.20.00752.

Bouchnak, I., and van Wijk, K.J.(2021). Structure, function, and substrates of Clp AAA+ protease systems in cyanobacteria, plastids, and apicoplasts: A comparative analysis. J Biol Chem 296:100338. 10.1016/j.jbc.2021.100338.

Dogra, V., Li, M., Singh, S., Li, M., and Kim, C.(2019). Oxidative post-translational modification of EXECUTER1 is required for singlet oxygen sensing in plastids. Nat Commun 10:2834. 10.1038/s41467-019-10760-6.

Dogra, V., Singh, R.M., Li, M., Li, M., Singh, S., and Kim, C.(2021). EXECUTER2 modulates the EXECUTER1 signalosome through its singlet oxygen-dependent oxidation. Mol Plant 10.1016/j.molp.2021.12.016.

Erbse, A., Dougan, D.A., and Bukau, B.(2003). A folding machine for many but a master of none. Nat Struct Biol 10:84–86.

Fu, Y., Li, X., Fan, B., Zhu, C., and Chen, Z.(2022). Chloroplasts Protein Quality Control and Turnover: A Multitude of Mechanisms. Int J Mol Sci 23 10.3390/ijms23147760.

Gao, L.L., Hong, Z.H., Wang, Y., and Wu, G.Z.(2023). Chloroplast proteostasis: A story of birth, life, and death. Plant Commun 4:100424. 10.1016/j.xplc.2022.100424.

Gibbs, D.J., Conde, J.V., Berckhan, S., Prasad, G., Mendiondo, G.M., and Holdsworth, M.J.(2015). Group VII Ethylene Response Factors Coordinate Oxygen and Nitric Oxide Signal Transduction and Stress Responses in Plants. Plant Physiol 169:23–31. 10.1104/pp.15.00338.

Hammarlund, E.U., Flashman, E., Mohlin, S., and Licausi, F.(2020). Oxygen-sensing mechanisms across eukaryotic kingdoms and their roles in complex multicellularity. Science 370 10.1126/science.aba3512.

Hanson, P.I., and Whiteheart, S.W.(2005). AAA+ proteins: have engine, will work. Nat Rev Mol Cell Biol 6:519–529. 10.1038/nrm1684.

Izumi, M., and Nakamura, S.(2018). Chloroplast Protein Turnover: The Influence of Extraplastidic Processes, Including Autophagy. Int J Mol Sci 19:15. 10.3390/ijms19030828.

Kim, J., Olinares, P.D., Oh, S.H., Ghisaura, S., Poliakov, A., Ponnala, L., and van Wijk, K.J.(2013). Modified Clp protease complex in the ClpP3 null mutant and consequences for chloroplast development and function in Arabidopsis. Plant Physiol 162:157–179. 10.1104/pp.113.215699.

Kim, J., Kimber, M.S., Nishimura, K., Friso, G., Schultz, L., Ponnala, L., and van Wijk, K.J.(2015). Structures, Functions, and Interactions of ClpT1 and ClpT2 in the Clp Protease System of Arabidopsis Chloroplasts. Plant Cell 27:1477-1496. 10.1105/tpc.15.00106.

Kohler, R.H., Cao, J., Zipfel, W.R., Webb, W.W., and Hanson, M.R.(1997). Exchange of protein molecules through connections between higher plant plastids. Science 276:2039–2042.

Lichtenthaler, H.K.(1987). Chlorophylls and Carotenoids: Pigments of photosynthetic Biomembranes. Methods in Enzymology 148:350–382.

Llamas, E., and Pulido, P.(2022). A proteostasis network safeguards the chloroplast proteome. Essays Biochem 66:219–228. 10.1042/EBC20210058.

Lupas, A.N., and Martin, J.(2002). AAA proteins. Curr Opin Struct Biol 12:746–753.

Majsec, K., Bhuiyan, N.H., Sun, Q., Kumari, S., Kumar, V., Ware, D., and van Wijk, K.J.(2017). The Plastid and Mitochondrial Peptidase Network in Arabidopsis thaliana: A Foundation for Testing Genetic Interactions and Functions in Organellar Proteostasis. Plant Cell 29:2687–2710. 10.1105/tpc.17.00481.

Montandon, C., Friso, G., Liao, J.R., Choi, J., and van Wijk, K.J.(2019). In Vivo Trapping of Proteins Interacting with the Chloroplast CLPC1 Chaperone: Potential Substrates and Adaptors. J Proteome Res 18:2585–2600. 10.1021/acs.jproteome.9b00112.

Nishimura, K., and van Wijk, K.J.(2015). Organization, function and substrates of the essential Clp protease system in plastids. Biochim Biophys Acta 1847:915–930. 10.1016/j.bbabio.2014.11.012.

Nishimura, K., Kato, Y., and Sakamoto, W.(2016). Chloroplast Proteases: Updates on Proteolysis within and across Suborganellar Compartments. Plant Physiol 171:2280–2293. 10.1104/pp.16.00330.

Nishimura, K., Apitz, J., Friso, G., Kim, J., Ponnala, L., Grimm, B., and van Wijk, K.J.(2015). Discovery of a Unique Clp Component, ClpF, in Chloroplasts: A Proposed Binary ClpF-ClpS1 Adaptor Complex Functions in Substrate Recognition and Delivery. Plant Cell 27:2677-2691. 10.1105/tpc.15.00574.

Nishimura, K., Asakura, Y., Friso, G., Kim, J., Oh, S.H., Rutschow, H., Ponnala, L., and van Wijk, K.J.(2013). ClpS1 is a conserved substrate selector for the chloroplast Clp protease system in Arabidopsis. Plant Cell 25:2276–2301. 10.1105/tpc.113.112557.

Obayashi, T., Hibara, H., Kagaya, Y., Aoki, Y., and Kinoshita, K.(2022). ATTED-II v11: A Plant Gene Coexpression Database Using a Sample Balancing Technique by Subagging of Principal Components. Plant Cell Physiol 63:869–881. 10.1093/pcp/pcac041.

Olivares, A.O., Baker, T.A., and Sauer, R.T.(2018). Mechanical Protein Unfolding and Degradation. Annu Rev Physiol 80:413–429. 10.1146/annurev-physiol-021317-121303.

Pulido, P., Llamas, E., Llorente, B., Ventura, S., Wright, L.P., and Rodriguez-Concepcion, M.(2016). Specific Hsp100 Chaperones Determine the Fate of the First Enzyme of the Plastidial Isoprenoid Pathway for Either Refolding or Degradation by the Stromal Clp Protease in Arabidopsis. PLoS Genet 12:e1005824. 10.1371/journal.pgen.1005824.

Rei Liao, J.Y., and van Wijk, K.J. (2019). Discovery of AAA+ Protease Substrates through Trapping Approaches. Trends Biochem Sci 44:528–545. 10.1016/j.tibs.2018.12.006.

Rei Liao, J.Y., Friso, G., Forsythe, E.S., Michel, E.J.S., Williams, A.M., Boguraev, S.S., Ponnala, L., Sloan, D.B., and van Wijk, K.J. (2022). Proteomics, phylogenetics, and coexpression analyses indicate novel interactions in the plastid CLP chaperone-protease system. J Biol Chem 298:101609. 10.1016/j.jbc.2022.101609.

Richter, A.S., Banse, C., and Grimm, B.(2019). The GluTR-binding protein is the heme-binding factor for feedback control of glutamyl-tRNA reductase. Elife 8 10.7554/eLife.46300.

Richter, A.S., Nagele, T., Grimm, B., Kaufmann, K., Schroda, M., Leister, D., and Kleine, T.(2023). Retrograde signaling in plants: A critical review focusing on the GUN pathway and beyond. Plant Commun 4:100511. 10.1016/j.xplc.2022.100511.

Rodriguez-Concepcion, M., D’Andrea, L., and Pulido, P.(2019). Control of plastidial metabolism by the Clp protease complex. J Exp Bot 70:2049–2058. 10.1093/jxb/ery441.

Rudella, A., Friso, G., Alonso, J.M., Ecker, J.R., and van Wijk, K.J.(2006). Downregulation of ClpR2 Leads to Reduced Accumulation of the ClpPRS Protease Complex and Defects in Chloroplast Biogenesis in Arabidopsis. Plant Cell 18:1704–1721.

Shimizu, T., and Masuda, T.(2021). The Role of Tetrapyrrole- and GUN1-Dependent Signaling on Chloroplast Biogenesis. Plants (Basel) 10 10.3390/plants10020196.

Su, T., Zhang, X.F., and Wu, G.Z.(2024). Functional conservation of GENOMES UNCOUPLED1 in plastid-to-nucleus retrograde signaling in tomato. Plant Sci:112053. 10.1016/j.plantsci.2024.112053.

Tadini, L., Jeran, N., and Pesaresi, P.(2020). GUN1 and Plastid RNA Metabolism: Learning from Genetics. Cells 9 10.3390/cells9102307.

Tapken, W., Kim, J., Nishimura, K., van Wijk, K.J., and Pilon, M.(2015). The Clp protease system is required for copper ion-dependent turnover of the PAA2/HMA8 copper transporter in chloroplasts. New Phytol 205:511–517. 10.1111/nph.13093.

van Dongen, J.T., and Licausi, F.(2015). Oxygen sensing and signaling. Annu Rev Plant Biol 66:345–367. 10.1146/annurev-arplant-043014-114813.

van Wijk, K.J., Leppert, T., Sun, Z., Kearly, A., Li, M., Mendoza, L., Guzchenko, I., Debley, E., Sauermann, G., Routray, P., et al.(2024). Detection of the Arabidopsis Proteome and Its Post-translational Modifications and the Nature of the Unobserved (Dark) Proteome in PeptideAtlas. J Proteome Res 23:185–214. 10.1021/acs.jproteome.3c00536.

Weits, D.A., van Dongen, J.T., and Licausi, F.(2021). Molecular oxygen as a signaling component in plant development. New Phytol 229:24–35. 10.1111/nph.16424.

Welsch, R., Zhou, X., Yuan, H., Alvarez, D., Sun, T., Schlossarek, D., Yang, Y., Shen, G., Zhang, H., Rodriguez-Concepcion, M., et al.(2018). Clp Protease and OR Directly Control the Proteostasis of Phytoene Synthase, the Crucial Enzyme for Carotenoid Biosynthesis in Arabidopsis. Mol Plant 11:149–162. 10.1016/j.molp.2017.11.003.

White, M.D., Klecker, M., Hopkinson, R.J., Weits, D.A., Mueller, C., Naumann, C., O’Neill, R., Wickens, J., Yang, J., Brooks-Bartlett, J.C., et al.(2017). Plant cysteine oxidases are dioxygenases that directly enable arginyl transferase-catalysed arginylation of N-end rule targets. Nat Commun 8:14690. 10.1038/ncomms14690.

Woodson, J.D.(2022). Control of chloroplast degradation and cell death in response to stress. Trends Biochem Sci 47:851–864. 10.1016/j.tibs.2022.03.010.

Wu, G.Z., and Bock, R.(2021). GUN control in retrograde signaling: How GENOMES UNCOUPLED proteins adjust nuclear gene expression to plastid biogenesis. Plant Cell 33:457–474. 10.1093/plcell/koaa048.

Wu, G.Z., Meyer, E.H., Wu, S., and Bock, R.(2019). Extensive Posttranscriptional Regulation of Nuclear Gene Expression by Plastid Retrograde Signals. Plant Physiol 180:2034–2048. 10.1104/pp.19.00421.

Wu, G.Z., Chalvin, C., Hoelscher, M., Meyer, E.H., Wu, X.N., and Bock, R.(2018). Control of Retrograde Signaling by Rapid Turnover of GENOMES UNCOUPLED1. Plant Physiol 176:2472–2495. 10.1104/pp.18.00009.

Wu, W., Zhu, Y., Ma, Z., Sun, Y., Quan, Q., Li, P., Hu, P., Shi, T., Lo, C., Chu, I.K., et al.(2013). Proteomic evidence for genetic epistasis: ClpR4 mutations switch leaf variegation to virescence in Arabidopsis. Plant J 76:943–956. 10.1111/tpj.12344.

Zhang, S., Zhang, H., Xia, Y., and Xiong, L.(2018). The caseinolytic protease complex component CLPC1 in Arabidopsis maintains proteome and RNA homeostasis in chloroplasts. BMC Plant Biol 18:192. 10.1186/s12870-018-1396-0.

Zybailov, B., Friso, G., Kim, J., Rudella, A., Rodriguez, V.R., Asakura, Y., Sun, Q., and van Wijk, K.J.(2009). Large scale comparative proteomics of a chloroplast Clp protease mutant reveals folding stress, altered protein homeostasis, and feedback regulation of metabolism. Mol Cell Proteomics 8:1789–1810.

